# A Cell Cycle-Dependent Ferroptosis Sensitivity Switch Governed by EMP2

**DOI:** 10.1101/2023.07.19.549715

**Authors:** Jason Rodencal, Nathan Kim, Veronica L. Li, Andrew He, Mike Lange, Jianping He, Amy Tarangelo, Zachary T. Schafer, James A. Olzmann, Julien Sage, Jonathan Z. Long, Scott J. Dixon

## Abstract

Ferroptosis is a non-apoptotic form of cell death characterized by iron-dependent lipid peroxidation. Ferroptosis can be induced by system x_c_^-^ cystine/glutamate antiporter inhibition or by direct inhibition of the phospholipid hydroperoxidase glutathione peroxidase 4 (GPX4). The regulation of ferroptosis in response to system x_c_^-^ inhibition versus direct GPX4 inhibition may be distinct. Here, we show that cell cycle arrest enhances sensitivity to ferroptosis triggered by GPX4 inhibition but not system x_c_^-^ inhibition. Arrested cells have increased levels of oxidizable polyunsaturated fatty acid-containing phospholipids, which drives sensitivity to GPX4 inhibition. Epithelial membrane protein 2 (EMP2) expression is reduced upon cell cycle arrest and is sufficient to enhance ferroptosis in response to direct GPX4 inhibition. An orally bioavailable GPX4 inhibitor increased markers of ferroptotic lipid peroxidation in vivo in combination with a cell cycle arresting agent. Thus, responses to different ferroptosis-inducing stimuli can be regulated by cell cycle state.

## INTRODUCTION

Ferroptosis is a non-apoptotic cell death mechanism that may be exploitable for cancer therapy.^1, 2^ Ferroptosis can be triggered by inhibition of the plasma membrane cystine/glutamate antiporter system x_c_^-^ or the glutathione-dependent lipid hydroperoxidase glutathione peroxidase 4 (GPX4).^3–5^ Small molecules inhibitors of system x_c_^-^ (e.g., erastin) and GPX4 (e.g., RSL3 and ML210) cause the lethal accumulation of membrane phospholipid hydroperoxides.^6–10^ However, cystine deprivation and direct GPX4 inhibition cause ferroptosis through related yet distinct mechanisms.^11^ For example, the polyunsaturated fatty acid (PUFA) metabolic enzyme acyl-CoA synthetase long chain family member 4 (ACSL4) may be more important for ferroptosis caused by direct GPX4 inhibition than by system x_c_^-^ inhibition or other stimuli.^11–13^ Molecular mechanisms that govern these distinct forms of ferroptosis remain poorly understood.

p53 is a transcription factor that can regulate diverse targets involved in cell cycle regulation, cell death, metabolism and other processes.^14, 15^ We and others find that stabilization of wild-type p53 in human cancer cells can promote resistance to ferroptosis triggered by system xc-inhibition.^16–18^ Mechanistically, this phenotype can involve altered protease function and the conservation of intracellular glutathione. Recent evidence suggests that p53 and canonical p53 target genes like CDKN1A (p21) may also influence GPX4 inhibitor sensitivity.^18–20^ However, published results are contradictory and the molecular mechanisms linking p53, CDKN1A, and potentially the cell cycle to altered ferroptosis sensitivity are unclear.

Here, we investigate how wild-type p53 expression impacts ferroptosis sensitivity. We find that wild-type p53 expression has two distinct effects on ferroptosis: suppression of ferroptosis in response to system x_c_^-^ inhibition and sensitization to ferroptosis in response to direct GPX4 inhibitors. We link sensitization to GPX4 inhibitors to cyclin dependent kinase (CDK) inhibition, cell cycle arrest, reduced expression of the tetraspanin protein EMP2, and altered levels of oxidizable polyunsaturated phosphatidylethanolamine (PUFA-PE) plasmalogens. We show that cell cycle arrest combined with GPX4 inhibition may enhance tumor cell lipid peroxidation in vivo. These results uncover a mechanistic link between the cell cycle and ferroptosis sensitivity via EMP2-dependent changes in lipid metabolism.

## RESULTS

### A screen for modulators of cell death upon p53 expression

Stabilization of wild-type p53 can enhance resistance to ferroptosis caused by system x_c_^-^ inhibition.^16–18^ We were initially curious whether p53-stabilized cells responded in a similar manner to other lethal stimuli and designed a small molecule screen to examine this question (Figure 1A). Wild-type p53 accumulates after treatment with the MDM2 inhibitor nutlin-3 (10 µM, 48 h)^16^. Because p53 expression itself can arrest cell proliferation, in this and all other experiments cells were seeded initially at densities that ensured a similar number of cells in both the control and nutlin-3-treated populations after 48 h. We then exposed these cells to 261 mechanistically diverse compounds, including kinase inhibitors, epigenetic modulators, and known inducers of different forms of cell death^8^, and measured cell death using the scalable time-lapse analysis of cell death kinetics (STACK) imaging method.^21^ The lethality of each compound was compared between cells pretreated with nultin-3 versus DMSO (Figure 1A).

**Figure 1.**
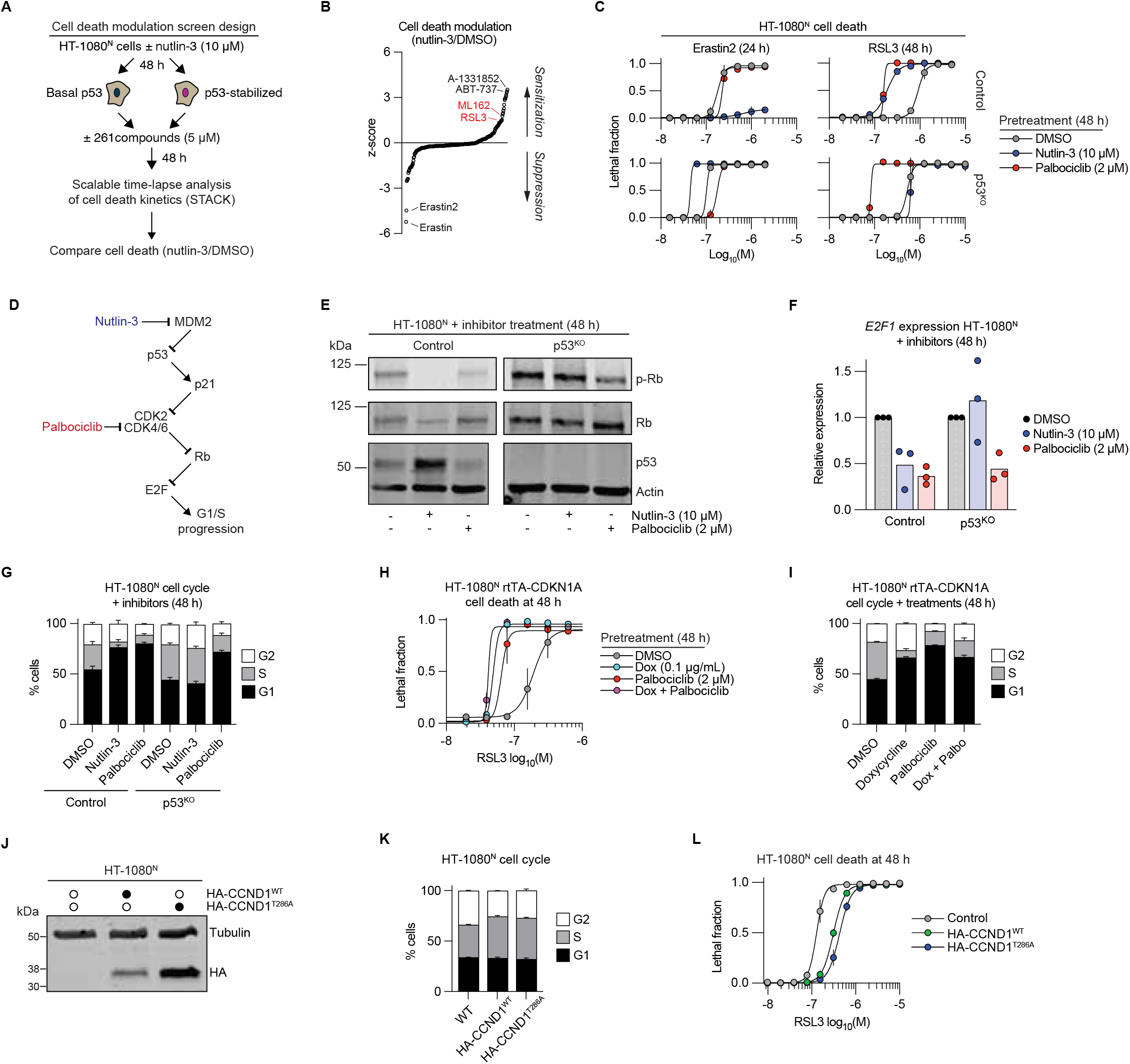
p53 stabilization sensitizes to GPX4 inhibitors. (A) Outline of the small molecule phenotypic screen. (B) Cell death modulation z-scores for HT-1080^N^ cells pretreated ± nutlin-3 (10 μM, 48 h) then treated ± 261 bioactive compounds (5 µM). The screen was repeated three times and z-scored mean values are shown. Compounds of interest are highlighted. (C) Cell death measured by imaging following pretreatment. (D) Outline of the CDK-Rb-E2F cell cycle pathway and associated inhibitors. (E) Protein abundance determined by immunoblotting. Blots are representative of three experiments. (F) Relative mRNA expression determined by reverse transcription and quantitative polymerase chain reaction (RT-qPCR) analysis. Individual datapoints from independent experiments are shown. (G) Cell cycle phase determined by propidium iodide (PI) staining and flow cytometry. Nutlin-3 was used at 10 µM and palbociclib at 2 μM. (H) Cell death measured by imaging following pretreatment. Dox, doxycycline. (I) Cell cycle phase determined by PI staining and flow cytometry. Dox was used at 0.1 μg/mL, palbociclib (Palbo) at 2 μM. (J) Protein abundance determined by immunoblotting. Blot is representative of three experiments. (K) Cell cycle phase determined by PI staining and flow cytometry in stable cell lines expressing the indicated constructs. (L) Cell death measured by imaging in stable cell lines expressing the indicated constructs. Data in (C), (G-I), (K) and (L) represent mean ± SD from three independent experiments.

As observed previously,^16–18^ p53 stabilization conferred resistance to the structurally similar small molecule system x_c_^-^ inhibitors erastin and erastin2, a result confirmed in focused assays using the same conditions (Figures 1B, 1C, and Table S1). Also as expected, p53 stabilization resulted in greater sensitivity to the pro-apoptotic BCL2/BCL-xL inhibitors ABT-737 and A-1331852 (Figure 1B).^22^ Unexpectedly, p53-stabilized cells were also sensitized to the covalent GPX4 inhibitors (GPX4i) RSL3 and ML162, a result we confirmed in focused assays (Figures 1B and 1C). Stabilization of wild-type p53 also reduced sensitivity to erastin2 while increasing sensitivity to RSL3 in Caki-1 renal carcinoma and A375 melanoma cells, consistent with a generalizable effect (Figures S1A and S1B). By contrast, decreased sensitivity to erastin2 and increased sensitivity to RSL3 were not observed in an established^16^ HT-1080 p53 gene disrupted (i.e., p53^KO^) cell line pretreated for 48 h with nutlin-3 (Figure 1C). Consistent with a p53-independent role for MDM2 in ferroptosis regulation,^18, 23^ nutlin-3 pre-treatment enhanced erastin2 sensitivity in HT-1080 p53^KO^ cells (Figure 1C), but this weak effect was not investigated further. Collectively, these findings indicated that p53 stabilization could confer resistance to ferroptosis triggered by system x_c_^-^ inhibition and simultaneously enhance sensitivity to ferroptosis caused by direct GPX4 inhibition.

### Cell cycle arrest sensitizes to GPX4 inhibition

Wild-type p53 can suppress ferroptosis induced by system x_c_^-^ inhibition by enhancing glutathione levels or by modulating other targets.^16, 17, 24^ How p53 expression sensitized to small molecule GPX4i was less clear, prompting us to investigate further. p53 has numerous transcriptional targets, including *CDKN1A*. CDKN1A (also known as p21) is an important regulator of CDK2, CDK4 and CDK6 that control G1/S progression via the retinoblastoma (Figure 1D).^25^ We tested the hypothesis that CDK4/6 inhibition was sufficient to sensitize to GPX4i by pretreating Control and p53^KO^ HT-1080^N^ cells with the CDK4/6 inhibitor palbociclib (2 µM, 48 h).^26^ Strikingly, both Control and p53^KO^ cells pretreated with palbociclib for 48 h were sensitized to RSL3 but not erastin2 (Figure 1C). Palbociclib treatment inhibited cell cycle progression in both Control and p53^KO^ cells as judged by reduced Rb phosphorylation, reduced *E2F1* expression, and cell cycle arrest at the G1/S transition, while nutlin-3 had these effects only in Control cells (Figures 1E-1G). Palbociclib pretreatment likewise reduced the expression of Rb-regulated cell cycle genes (*E2F1*, *CCNA2*) and selectively sensitized to RSL3 but not erastin2 in Caki-1 and A375 cells, without altering p53 levels (Figures S1A-S1F).

We further examined the relationship between proliferative arrest and ferroptosis sensitivity. In HT-1080^N^ cells, the duration of palbociclib (2 µM) pretreatment correlated with increased RSL3 potency (Figure S2A). The concentration of palbociclib used in the pretreatment also positively correlated with RSL3 potency (Figure S2B). In these cells, RSL3 potency was also tightly correlated with the degree of proliferative arrest (Pearson R^2^ = 0.95, p < 0.001) (Figure S2C). Pretreatment with the structurally distinct CDK4/6 inhibitor abemaciclib likewise reduced *E2F1* expression, slowed cell cycle progression and sensitized HT-1080^N^ cells to RSL3 (Figures S2D-S2F). Similar results were also obtained using doxycycline-induced overexpression of CDKN1A (Ref.^16^), which arrested cells mostly in G1, reduced *E2F1* expression, and sensitized to GPX4i treatment (Figures 1H, 1I, S2G, and S2H). Notably, CDKN1A overexpression and palbociclib treatment did not have additive effects on RSL3 sensitivity, suggesting they act through the same mechanism (Figure 1H).

To further extend these results we employed complementary gain of function approaches to test whether increased cell cycle function reduced cell death sensitivity. Towards this end, we stably overexpressed the CDK4/6 subunit CCND1 (cyclin D) in HT-1080^N^ cells via lentiviral transduction. CCND1 overexpression was sufficient to enhance the expression of cell cycle-associated genes (*E2F1*, *CCNA2*), increase the number of cells in S phase, and reduce sensitivity to RSL3 (Figure 1J-1L, and S2I). Expression of a degradation-resistant CCND1^T286A^ mutant, which accumulates to higher levels than the wild-type protein, did not further increase the number of S-phase cells or further reduce ferroptosis sensitivity (Figures 1J-1L, and S2I). Collectively, these results indicated that cell cycle state can alter sensitivity to GPX4i-induced ferroptosis.

### Cell cycle arrest sensitizes to lipid peroxidation upon GPX4 inhibition

We hypothesized that CDK4/6 inhibition sensitized cells to GPX4i-induced ferroptosis and not a different lethal mechanism. Consistent with this hypothesis, GPX4i-induced cell death in both vehicle (DMSO)-pretreated and palbociclib-pretreated HT-1080^N^ cells was fully inhibited by the canonical small molecule ferroptosis inhibitors ferrostatin-1 and deferoxamine, but not the pan-caspase inhibitor Q-VD-OPh (Figures 2A and S3A). Thus, palbocilcib pretreatment sensitized to ferroptosis and not some other cell death mechanism.

**Figure 2.**
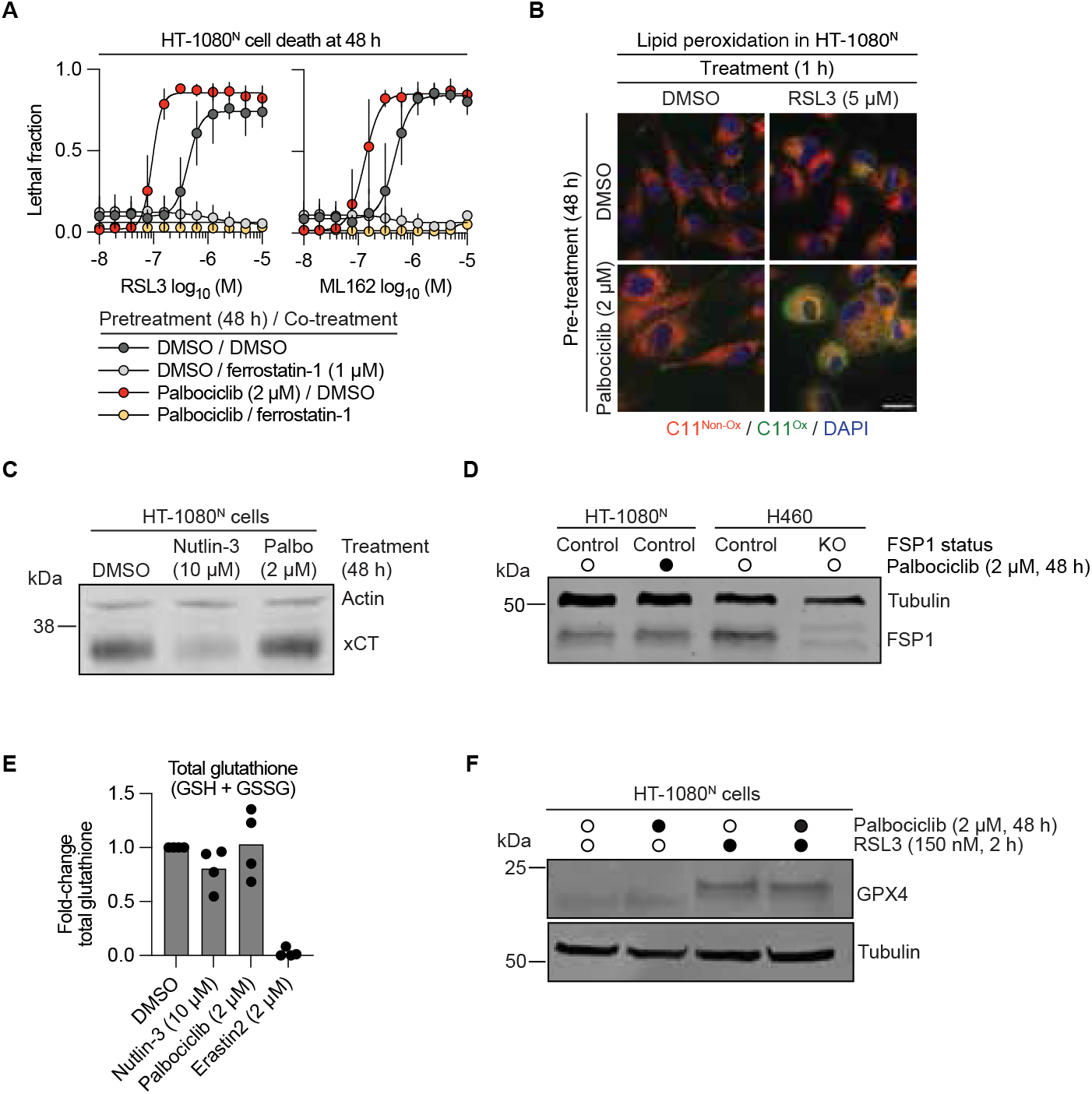
GPX4 inhibitor sensitization involves increased lipid peroxidation. (A) Cell death measured by imaging following pretreatment. Data represent mean ± SD from three independent experiments. (B) C11 BODIPY 581/591 (C11) imaging by confocal microscopy. Non-ox: non-oxidized, Ox: oxidized. Scale bar = 20 µm. Images are representative of three experiments. (C) Protein abundance determined by immunoblotting. Palbo, palbociclib. (D) Protein abundance determined by immunoblotting. H460 Control and *FSP1* gene-disrupted (KO) cell lines are included as an anti-FSP1 antibody control. (E) Relative total glutathione levels determined using Ellman’s reagent. Nutlin-3 and palbociclib treatments were for 48 h. Erastin2 treatment (8 h) is a positive control for glutathione depletion. Individual data points from four independent experiments are shown. (F) Protein abundance and migration determined by immunoblotting in one-dimensional SDS-PAGE gels. Cells were pretreated ± palbociclib for 48 h before treatment ± RSL3. Blots in (C), (D) and (F) are representative of three independent experiments.

Increased membrane lipid peroxidation is the defining feature of ferroptosis.^8, 10^ In HT-1080^N^ cells pretreated with palbociclib, we observed using confocal microscopy greater oxidation of the lipid peroxidation probe C11 BODIPY 581/591 (C11) in response to GPX4i treatment (Figure 2B). Doxycycline (Dox)-inducible CDKN1A expression in HT-1080^N^ cells likewise enhanced C11 oxidation in response to GPX4 inhibition, as detected using epifluorescent imaging (Figure S3B). We further extended these results to additional cell models, showing that 786-O renal cell carcinoma and MDA-MB-231 triple negative breast cancer cells had enhanced sensitivity to RSL3 and ML210 following palbociclib pretreatment along with increased sensitivity to C11 oxidation (Figures S3C and S3D). Thus, CDK4/6 inhibition enhanced GPX4i-induced membrane lipid peroxidation.

We initially hypothesized that enhanced lipid peroxidation and ferroptosis sensitivity were due to reduced levels of known anti-ferroptosis proteins or metabolites. However, palbociclib did not reduce the abundance of xCT (the antiporter subunit of system x_C_^-^), GPX4, or ferroptosis suppressor protein 1 (FSP1) protein, nor did it reduce total intracellular glutathione levels (Figure 2C-2E, and S3E), eliminating several plausible sensitization mechanisms.^1^ Another hypothesis was that CDK4/6 inhibition increased the susceptibility of GPX4 to inhibitor binding. We used one-dimensional SDS-PAGE to examine the apparent migration of GPX4, which is slowed by GPX4i binding.^27^ Palbociclib pretreatment did not alter the apparent migration of GPX4, either when used as a single agent or in combination with a low lethal dose of RSL3, suggesting that target engagement was unaffected (Figure 2F). Thus, sensitization to membrane lipid peroxidation and ferroptosis following CDK4/6 inhibition did not appear to be explained by altered interactions between RSL3 and GPX4.

### Cell cycle arrest is associated with phospholipid remodeling

We next considered whether proliferative arrest enhanced GPX4i sensitivity by altering membrane phospholipid (PL) composition. The balance between pro-ferroptotic PUFA-containing PLs and anti-ferroptotic MUFA-containing PLs is a key determinant of overall ferroptosis sensitivity^8, 28^. We envisioned that increased PUFA-PL levels and/or decreased MUFA-PL levels could underlie the greater sensitivity of arrested cancer cells to GPX4i. To test this hypothesis, we compared HT-1080^N^ Control to *ACSL4* gene-disrupted (“KO”) cell lines which have reduced levels of various PUFA-PLs and PUFA-containing triacylglycerols^8, 12^. Consistent with our hypothesis, palbociclib pretreatment did not sensitize ACSL4^KO1/2^ cells to RSL3 despite effective inhibition of Rb phosphorylation and reduced expression of *E2F1* (Figures 3A-3D). Thus, cell cycle arrest per se was insufficient to sensitize to GPX4i-induced ferroptosis in the absence of a pro-ferroptotic lipid metabolic environment.

**Figure 3.**
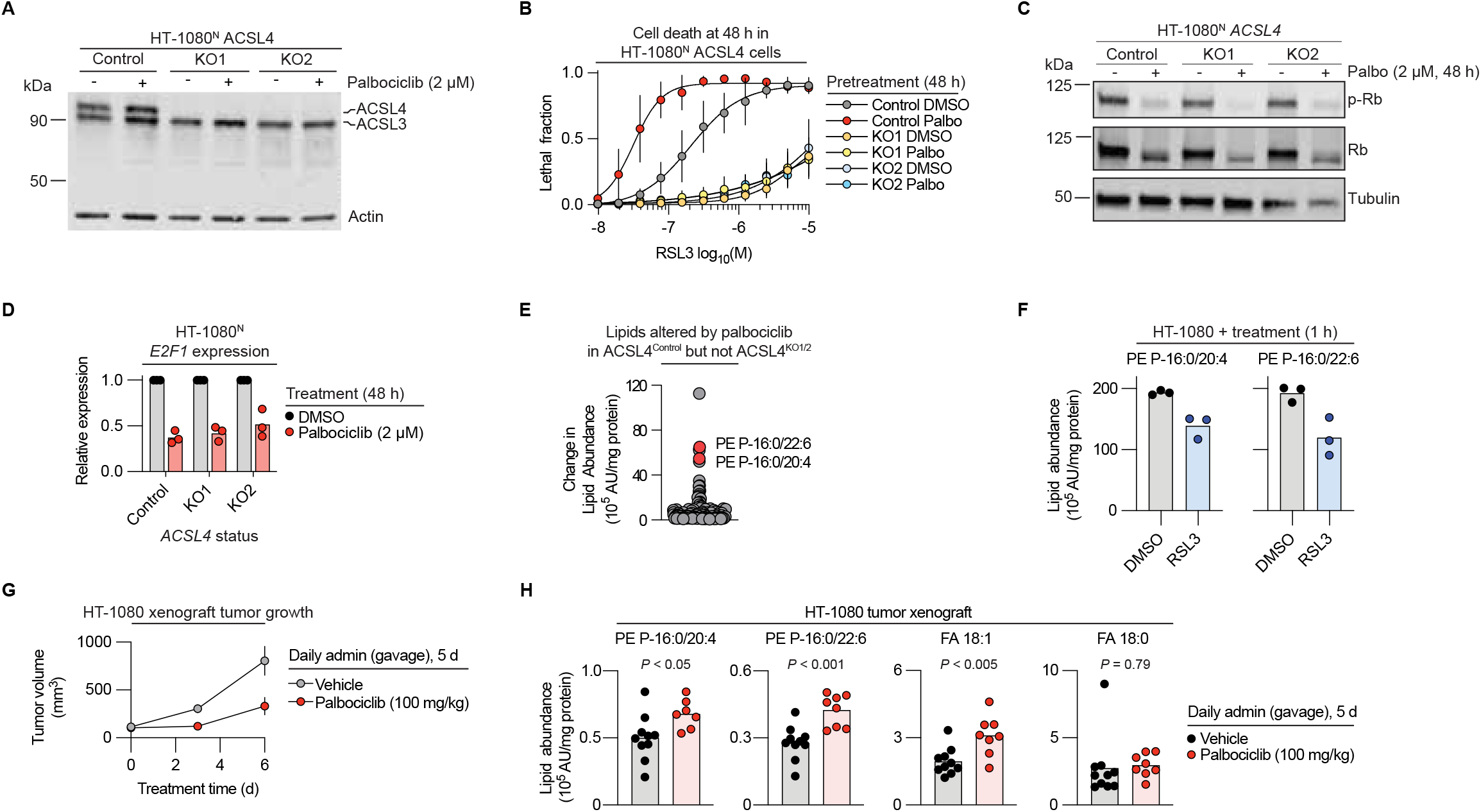
Altered lipid peroxidation sensitivity requires ACSL4. (A) Protein abundance determined by immunoblotting. KO1, gene-disrupted cell line 1. KO2, gene-disrupted cell line 2. Cell line genotypes and protein phenotypes have been reported previously.^8, 12^ (B) Cell death measured by imaging following pretreatment. Results represent mean ± SD from three independent experiments. (C) Protein abundance determined by immunoblotting. Palbo: palbociclib. (D) Relative *E2F1* expression determined by reverse transcription and quantitative polymerase chain reaction (RT-qPCR) analysis. (E) Differentially abundant (FDR *q* < 0.05) lipids following palbociclib (2 µM, 48 h) treatment in Control versus ACSL4^KO1/2^ cell lines, as determined by liquid chromatography coupled to mass spectrometry (LC-MS) analysis (see Materials and Methods). Each dot represents one species and is the average of five or six independent cultures examined in parallel. Lipids confirmed by MS/MS analysis are indicated. AU, arbitrary units. (F) Analysis of lipid abundance by LC-MS. RSL3 was used at 1 µM. AU, arbitrary units. (G) Analysis of HT-1080 tumor xenografts growth. Tumors were allowed to grow to an average size of ∼100 mm^3^, then mice were treated ± palbociclib daily for 5 days (tumors were harvested on day 6 for further analysis). Data represent mean ± SEM (n = 8 or 10 tumors per condition). (H) Lipid abundance following treatment, as determined by liquid chromatography coupled to mass spectrometry (LC-MS) analysis. Data from individual tumors is shown. Two-tailed Student’s t-test was used to compare mean differences between groups. AU, arbitrary units. Blots in (A) and (C) are representative of three experiments. Individual datapoints from separate experiments are shown in (D) and (F).

To understand which lipids were altered by cell cycle arrest, we performed untargeted lipidomic analysis using liquid chromatography coupled to mass spectrometry (LC-MS) in Control and ACSL4^KO1/2^ cells. We hypothesized that lipids altered by palbociclib in Control cells, but not ACSL4^KO1/2^ cells, would be candidate regulators of ferroptosis sensitivity. Two species that changed substantially in abundance were confirmed by MS/MS analysis as the PUFA-containing ether-linked phospholipids PE P-16:0/20:4 and PE P-16:0/22:6 (Figure 3E, Table S2). Interestingly, the abundances of both PE P-16:0/20:4 and PE P-16:0/22:6 were reduced following treatment with a lethal dose of RSL3 where cells were harvested just before the onset of cell death (Figure 3F). This suggested that that these two lipids could be converted to different species due to oxidation during ferroptosis.

Next, we examined whether changes observed in cultured cells in response to palbociclib were relevant in vivo. We generated HT-1080 xenograft tumors in the flanks of NOD-*scid* IL2Rgamma^null^ (NSG) mice and, once these tumors reached ∼100 mm^3^ in size, started treatment with palbociclib (100 mg/kg) for five days. As expected, palbociclib-treated tumors did not increase in size over the course of the experiment (Figure 3G). Moreover, in harvested tumors, we observed that palbociclib treatment significantly increased the levels of PE P-16:0/20:4 and PE P-16:0/22:6, consistent with results obtained in cultured cells (Figure 3H). We also observed an increase in the levels of the MUFA 18:1 free fatty acid, with no corresponding change in the levels of the fully saturated 18:0 precursor. These findings indicated that palbociclib treatment increased the levels of PUFA-containing PLs in cultured cells and in vivo, correlating with greater lethal lipid peroxidation and induction of ferroptosis in response to GPX4 inhibition.

### Identification of cell cycle-regulated genes linked to ferroptosis

CDK4/6 regulate the transcription of hundreds of genes via the Rb-E2F pathway.^29^ To identify genes that could underlie sensitization to GPX4i upon cell cycle arrest we integrated gene expression profiling with the analysis of publicly available gene expression and compound activity datasets. Using RNA sequencing (RNA-seq) we identified 629 genes whose expression was significantly altered by palbociclib treatment (FDR *q* < 0.05) (Figure 4A and 4B, Table S3). We hypothesized that other treatments that arrest cell proliferation through distinct means might alter gene expression and enhance GPX4i sensitivity in a similar manner. The mitogen activated protein kinase (MAPK) cascade regulates cell proliferation in part by promoting expression of CCND1.^30^ Consistent with our hypothesis, inhibition of MAPK signaling using the MAPK kinase (MEK1/2) inhibitor pimasertib (0.5 µM, 48 h) arrested cell proliferation and, like palbociclib, sensitized HT-1080^N^ cells to GPX4i treatment (Figures S4A-S4C). We therefore performed RNA-seq on pimasertib-treated cells and intersected the lists of genes whose expression was significantly altered by both palbociclib and pimasertib. The two datasets shared 498 genes in common, including canonical cell cycle genes (e.g., *MKI67*, *CCNA2*, see Table S3) that were downregulated in both conditions.^31^ Next, we focused on genes whose basal mRNA expression correlated significantly with sensitivity to at least two of three GPX4i (RSL3, ML210 or ML162) tested as part of Cancer Therapeutics Response Portal database,^32^ and that have likely E2F binding sites in their promoter regions based on chromatin immunoprecipitation (ChIP-seq) analysis (TargetGeneReg 2.0).^33^ This integrated analysis ultimately pinpointed four genes: *MBOAT1*, *GPD2, EMP2*, and *MYO19* (Figures 4A-4C).

**Figure 4.**
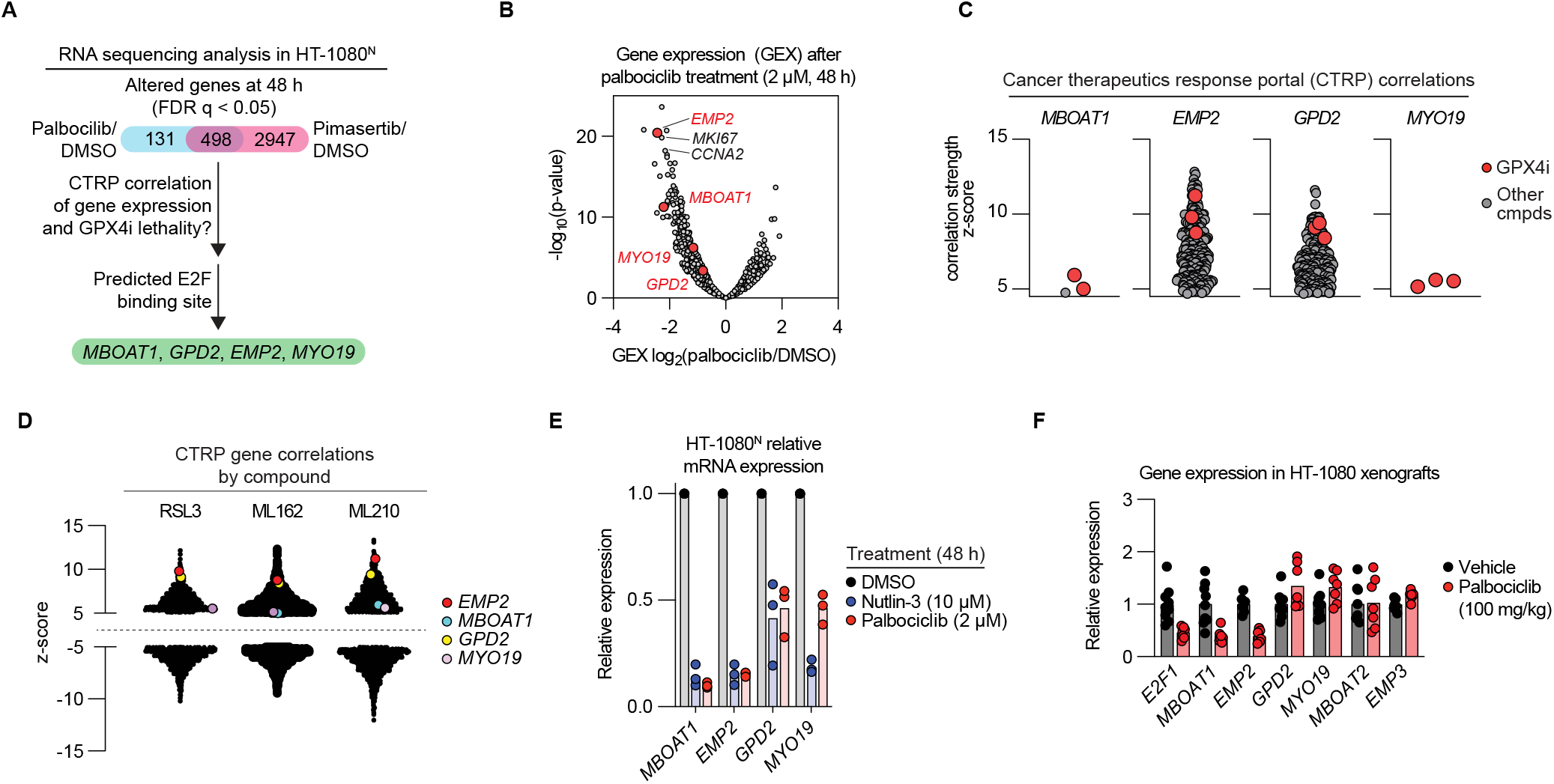
Identification of cell cycle-regulated genes associated with ferroptosis. (A) Overview of the pipeline used to identify candidate regulators of ferroptosis sensitivity. CTRP, cancer therapeutics response portal. (B) RNA-sequencing. Labeled red points are from (A) (C-D) Data from the CTRP database correlating compound sensitivity and basal gene expression. Points in (C) represent an individual compounds correlated with the labeled gene. Points in (D) represent individual genes correlated with the labeled compound. (E) Relative mRNA expression determined by reverse transcription and quantitative polymerase chain reaction (RT-qPCR) analysis. Individual datapoints from separate experiments are shown. (F) Relative mRNA expression determined by RT-qPCR analysis from tumor xenografts harvested previously. Individual datapoints represent separate xenograft tumors. Data is normalized to mean ΔΔCT values of vehicle control for each gene.

MBOAT1 (membrane bound O-acyltransferase domain containing 1) is a MUFA-specific lysophospholipid acyltransferase (LPLAT) recently linked to ferroptosis regulation.^34, 35^ Less oxidizable MUFAs can suppress ferroptosis by displacing more oxidizable PUFA species from plasma membrane phospholipids.^8, 36, 37^ Lower *MBOAT1* expression upon cell cycle arrest could sensitize to GPX4i by reducing the levels of anti-ferroptotic MUFA-PLs^8, 35, 38^ and/or allowing for increased levels of PUFA-PLs, consistent with our lipidomic results (Figure 3). GPD2 (glycerol-3-phosphate dehydrogenase 2) is a mitochondrial enzyme that helps generate reduced Coenzyme Q10 (CoQ10), a ferroptosis inhibitory metabolite.^39^ Downregulation of *GPD2* may explain sensitization to GPX4i. Connections between EMP2 (epithelial membrane protein 2) or MYO19 (myosin XIX) and ferroptosis were less obvious. However, reciprocal analysis of the CTRP dataset revealed that basal *EMP2* expression was among the strongest correlates of sensitivity to GPX4i across the entire transcriptome (Figure 4D).

We sought independent evidence that these four genes were likely to be cell cycle regulated. From a database of transcriptomic profiles acquired at various points across the cell cycle of cultured HeLa cells^40^ we observed that *MBOAT1* expression peaked at the G1/S transition (periodic rank: 178), similar to *E2F1* (periodic rank: 134), while *GPD2* expression was more dispersed across the cell cycle (periodic rank: 1,501). No data was available in this analysis for *EMP2* or *MYO19*. However, EMP2 overexpression can arrest lung cancer cells in G1, consistent with a link to cell cycle regulation.^41^

To investigate further, we performed targeted gene expression analysis in HT-1080^N^ cells. All four candidate genes were downregulated in HT-1080^N^ cells treated with nutlin-3 and/or palbociclib treatment, with *MBOAT1* and *EMP2* showing the strongest downregulation in cultured HT-1080 cells (Figure 4E). Strong transcriptional downregulation of *MBOAT1* and *EMP2*, but not *GPD2* or *MYO19*, was also observed in HT-1080 xenograft tumors treated with palbociclib (Figure 4F). Transcription of the closely related genes *MBOAT2* (to *MBOAT1*) and *EMP3* (to *EMP2*) were not decreased by palbociclib treatment (Figure 4F). Unfortunately, we were unable to examine effects at the protein level as we could not identify antibodies that detected endogenous MBOAT1 or EMP2 protein, a limitation of this study. Nonetheless, based on these results, we focused further on *MBOAT1* and *EMP2* as candidate cell cycle-dependent regulators of ferroptosis.

### MBOAT1 can regulate ferroptosis sensitivity

MBOAT1 uses oleoyl-CoA (a MUFA) as a substrate to synthesize MUFA-PLs (Figure 5A).^34, 35, 42^ MBOAT1 encodes one of four mammalian MBOAT-family lysophospholipid acyltransferases (MBOAT1, MBOAT2, MBOAT5/LPCAT3, and MBOAT7), each with partially distinct substrate specificities.^43^ Consistent with MBOAT1 regulation by the cell cycle, nutlin-3 reduced *MBOAT1* expression in a p53-dependent manner while palbociclib and pimasertib reduced *MBOAT1* expression in both HT-1080 Control and p53^KO^ cells (Figure 5B). *MBOAT2, MBOAT5/LPCAT3,* and *MBOAT7* are all classified as non-periodic genes in Cyclebase.^40^ Indeed, Dox-induced CDKN1A overexpression in HT-1080 cells reduced the expression of *MBOAT1* but not *MBOAT2* or *MBOAT5* (Figure 5C), and palbociclib treatment reduced *MBOAT1* expression in cultured A375^N^ (p53 wild-type), T98G^N^ (p53 mutant), and H1299^N^ (p53 null) cells while *MBOAT2* and/or *MBOAT5* expression was not reduced (Figure 5D). Thus, various cell cycle arresting conditions resulted in selective downregulation of *MBOAT1* expression.

**Figure 5.**
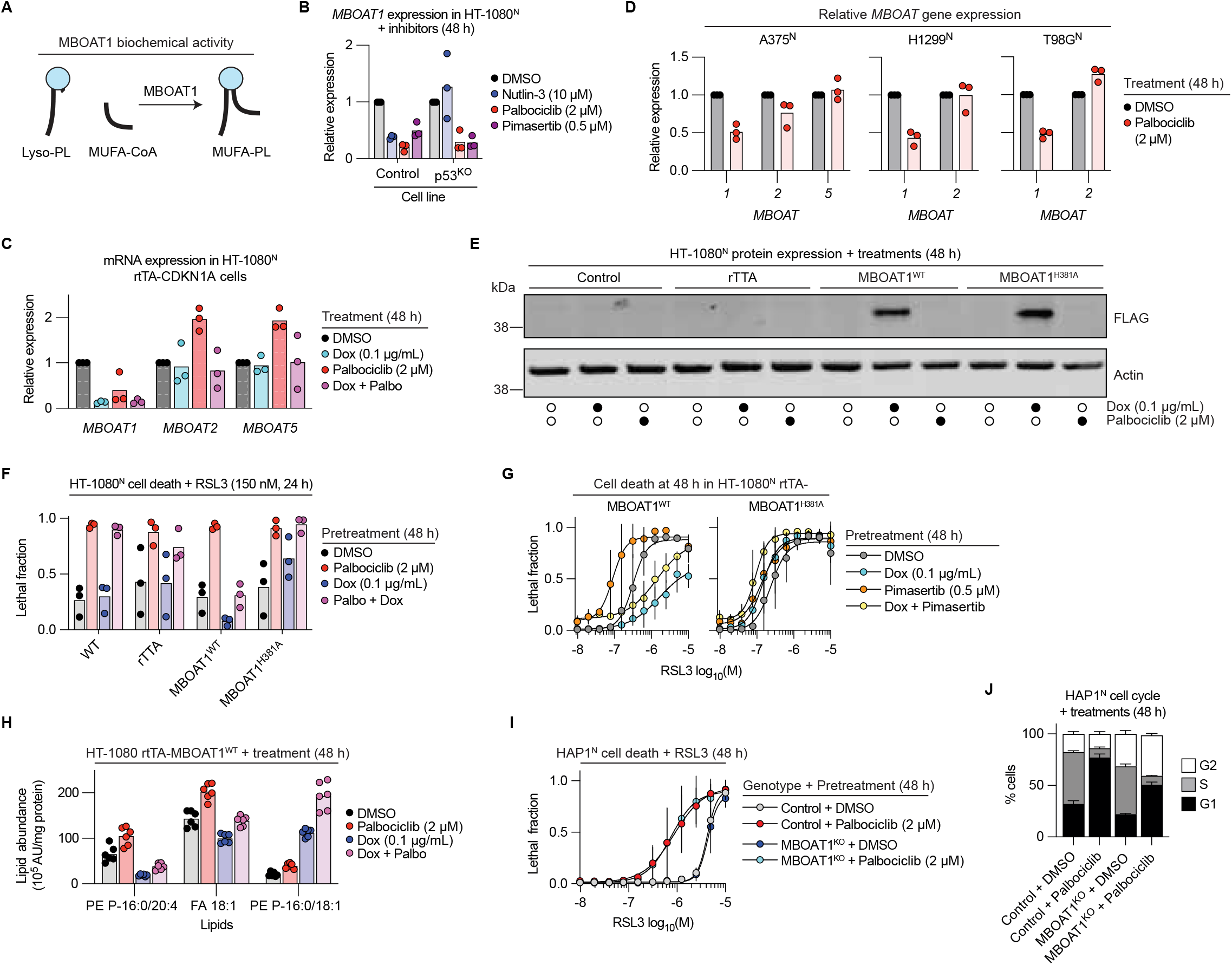
MBOAT1 can regulate ferroptosis sensitivity. (A) Cartoon schematic of MBOAT1 function. (B-D) Relative mRNA expression determined by reverse transcription and quantitative polymerase chain reaction (RT-qPCR) analysis. Dox, doxycycline. (E) Protein abundance determined by immunoblotting. Blots are representative of three experiments. (F,G) Cell death measured by imaging following pretreatment. (H) Lipid abundance following treatment, as determined by liquid chromatography coupled to mass spectrometry (LC-MS) analysis. AU, arbitrary units. (I) Cell death measured by imaging following pretreatment. (J) Cell cycle phase determined by propidium iodide (PI) staining and flow cytometry. Palbociclib was used at 2 μM. Individual datapoints from independent experiments are shown in (B-D), (F) and (H). Data in (G), (I) and (J) represent mean ± SD from three independent experiments.

To test whether reduced *MBOAT1* expression was necessary for enhanced sensitivity to GPX4i upon cell cycle arrest, we engineered HT-1080 cell lines where epitope (FLAG)-tagged wild-type MBOAT1 (MBOAT1^WT^) or a predicted catalytic-dead MBOAT1^H381A^ mutant^44^ were expressed under Dox-inducible control (Figure 5E). Dox-induced overexpression of MBOAT1^WT^ prevented palbociclib-mediated sensitization to RSL3 (Figure 5F). By contrast, overexpression of MBOAT1^H381A^ exacerbated the lethality of RSL3, possibly due to a dominant negative effect (Figure 5F). Similar results were obtained when using pimasertib pretreatment instead of palbociclib pretreatment, demonstrating that these effects were not limited to direct CDK4/6 inhibition (Figure 5G). Importantly, MBOAT1^WT^ overexpression did not prevent palbociclib from reducing Rb phosphorylation, lowering *E2F1* expression, or arresting cell cycle progression (Figures S5A-C). Collectively, these results demonstrated that *MBOAT1* overexpression was sufficient to revert sensitization to GPX4i upon cell cycle arrest.

We hypothesized that MBOAT1 expression would suppress ferroptosis by altering lipid metabolism. Consistent with this hypothesis, Dox-inducible overexpression of MBOAT1^WT^ but not MBOAT1^H381A^ reduced the levels of PE P-16:0/20:4 and prevented the accumulation of PE P-16:0/20:4 upon palbociclib treatment (Figure 5H). By contrast, the basal level of FA 18:1 (oleic acid) was decreased by MBOAT1^WT^ expression and was only partially recovered by palbociclib treatment. A different MUFA species, PE *a*-16:1/18:1, where ‘*a*’ denotes that this alkyl lipid may be either an ether lipid or plasmalogen, was largely unaffected by palbociclib treatment but was substantially increased upon MBOAT1^WT^ expression (Figure 5F). By contrast, Dox-induced MBOAT1^H381A^ expression did not result in similar changes in the abundance of these PUFA and MUFA species (Figure S5D). We inferred that MBOAT1 suppressed GPX4i sensitivity by reducing PUFA-PL levels and/or increasing MUFA-PL levels, consistent with recent findings.^35^

We next investigated whether *MBOAT1* loss of function alone was sufficient to enhance ferroptosis sensitivity and obtained evidence that this was not the case. Disruption of *MBOAT1* in human HAP1 haploid cells did not enhance sensitivity to RSL3 basally or alter the response of these cells to palbociclib pretreatment (Figure 5I). We confirmed that palbociclib did reduce the number of S-phase cells in MBOAT1^KO^ cells (Figure 5J). Of note, untreated HAP1 MBOAT1^KO^ cells displayed a greater propensity to arrest in the G2/M phase, consistent with some role for MBOAT1 in progression through the cell cycle, but this was not examined further here (Figure 5J). In line with the cell death results obtained in HAP1 cells, a short interfering RNA (siRNA) targeting *MBOAT1* did not enhance ferroptosis sensitivity in HT-1080^N^, T98G^N^, or A375^N^ cell lines (Figures S5E-S5J). Collectively, these findings indicated that while *MBOAT1* downregulation may be necessary for sensitization to GPX4i upon cell cycle arrest, reduced expression of *MBOAT1* alone was not sufficient to enhance GPX4i sensitivity in normally cycling cells.

### EMP2 downregulation sensitizes to ferroptosis

We next turned our attention to *EMP2*. Low basal EMP2 expression was highly correlated to all three GPX4i in the CTRP (Figure 4D), and also significantly downregulated in response to palbociclib in both cultured cells and tumor xenografts (Figures 4E and 4F). EMP2 is one of four related peripheral myelin protein 22-family tetraspanin genes: *PMP22*, *EMP1*, *EMP2*, and *EMP3*.^45^ Only *EMP2* was significantly downregulated by palbociclib or pimasertib in our RNA-Seq analysis (Table S3). We confirmed in HT-1080^N^ cells that palbociclib and abemaciclib both reduced *EMP2* expression with little effect on the expression of *PMP22* and *EMP3* and a weaker effect on *EMP1* (Figure 6A). Dox-inducible CDKN1A overexpression in HT-1080^N^ cells strongly reduced *EMP2* expression, and the effect of palbociclib was not additive with CDKN1A overexpression, consistent with a single pathway (Figure 6B). *EMP2* expression was also substantially downregulated in other p53 wildtype cells (Caki-1^N^, A375^N^) upon treatment with nutlin-3 and/or pimasertib (Figure S6A). Collectively, these findings were consistent with *EMP2* downregulation upon cell cycle arrest.

**Figure 6.**
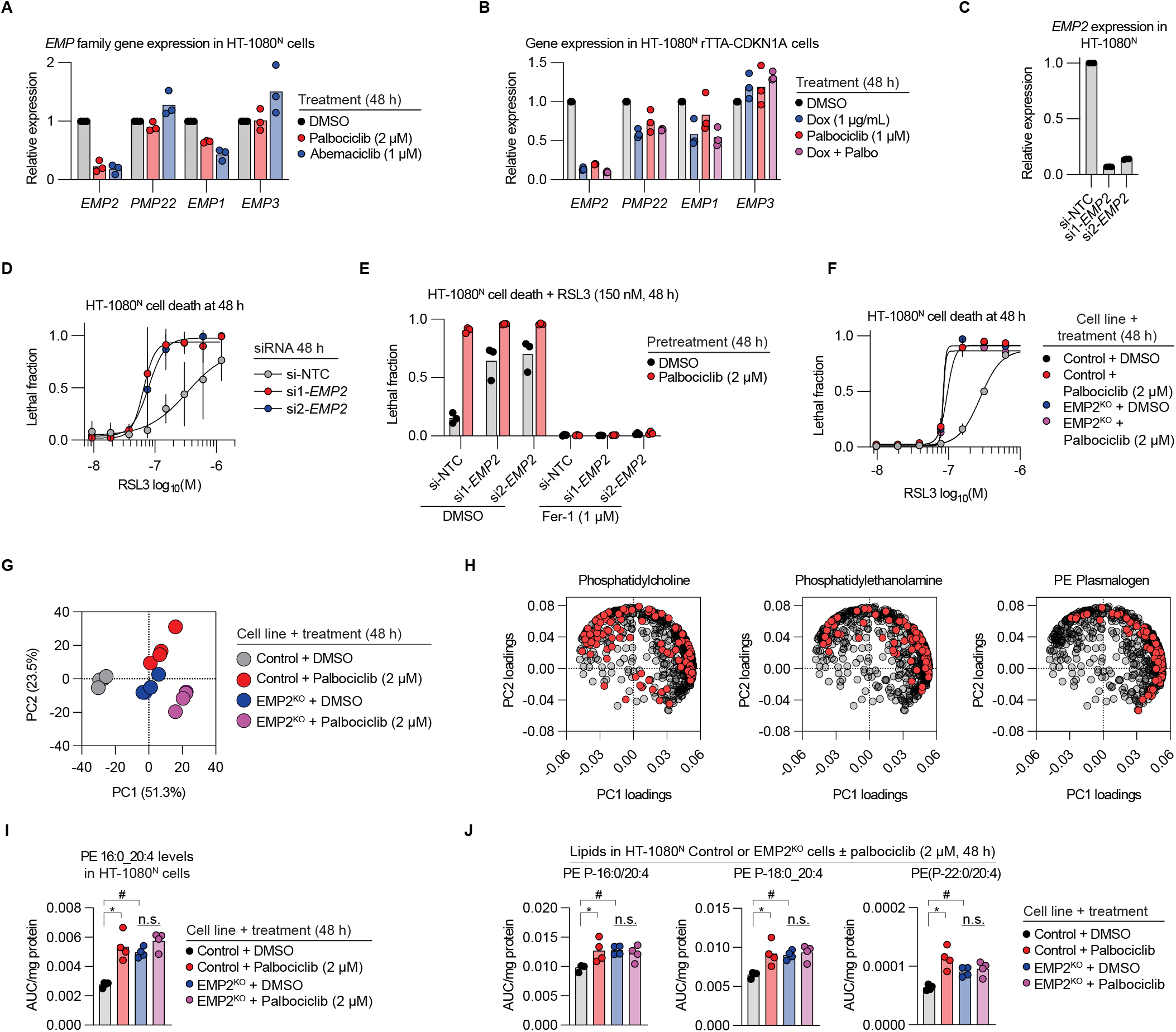
EMP2 regulates ferroptosis sensitivity. (A-C) Relative mRNA expression determined by reverse transcription and quantitative polymerase chain reaction (RT-qPCR) analysis. Dox, doxycycline. si, short interfering RNA. NTC, non-targeting control. (D-F) Cell death measured by imaging following pretreatment. Fer-1, ferrostatin-1. (G) Principal component analysis (PCA) scores plot. Each point represents one sample. (H) PCA loadings plots. Each red point belongs to the lipid class labeled in the graph title. (I-J) Lipid abundance following treatment, as determined by liquid chromatography coupled to mass spectrometry (LC-MS) analysis. *Indicates FDR q < 0.05. #Indicates FDR q < 0.01. AUC, area under the curve. Individual datapoints from three (A-C), (E), or four (I) and (J) independent experiments are shown. Results in (D), (F) represent mean ± SD from three independent experiments.

Given our experience with *MBOAT1*, we next examined whether loss of *EMP2* function alone was sufficient to regulate ferroptosis sensitivity. Strikingly, in HT-1080^N^ cells genetic silencing of *EMP2* alone using two different siRNAs was sufficient to sensitize to RSL3-induced ferroptosis, almost to the level of palbociclib alone (Figures 6C-E). This sensitization was due to the induction of ferroptosis, as it was fully suppressed by ferrostatin-1 (Figure 6E). Similar results were obtained using these siRNAs in Caki-1 cells (Figures S6B and S6C).

We next isolated a clonal HT-1080^N^ *EMP2* gene-disrupted cell line (EMP2^KO^) and a matched Control cell line that had undergone the CRISPR process without modification. Using genomic DNA sequencing, we confirmed that EMP2^KO^ cells contained an early stop codon (see Methods). Moreover, by RT-qPCR analysis we detected reduced *EMP2* expression in the gene-disrupted cell line and no change in the expression of *EMP3* (Figure S6D). Both the Control and EMP2^KO^ cell lines were sensitive to the effects of palbociclib, exhibiting equivalent reductions in the mRNA levels of the cell cycle genes *E2F1* and *CCNA2* (Figure S6E), and both cell lines arrested in G1 after palbociclib treatment (Figure S6F), indicating that EMP2 disruption per se did not alter the cell cycle or the ability of CDK4/6 inhibition to arrest cell cycle progression. We also showed that *EMP2* disruption did not alter *MBOAT1* transcript levels (Figure S6G). Compared to the Control cell line, EMP2^KO^ cells were strongly sensitized to the GPX4i RSL3 and ML210, more weakly sensitized to erastin2, and not sensitized to the pro-apoptotic agent bortezomib (Figure S6H). Consistent with results obtained using siRNA treatment, palbociclib treatment and genetic disruption of *EMP2* each sensitized to RSL3 treatment to an equivalent extent, and these effects were not additive, implying action through a common mechanism (Figure 6F).

### Loss of *EMP2* promotes PUFA enrichment in phospholipids

Palbociclib treatment causes the accumulation of PUFA-PEs in vitro and in vivo (Figures 4E, and 4H). We hypothesized that loss of *EMP2* may sensitize to ferroptosis by promoting the accumulation of PUFA-PEs. Using untargeted LC-MS/MS, we detected 678 lipid species with high confidence in Control and EMP2^KO^ cells treated with DMSO or palbociclib (Figure S7A). Given that loss of *EMP2* is sufficient to promote GPX4i sensitization, and that functional *EMP2* is necessary for palbociclib to sensitize to GPX4i (Figure 6F), we designed a strategy to filter our initial list to focus on lipids that may be relevant for this phenotype (Figure S7A and Methods). While saturated fatty acid (SFA), MUFA, and PUFA-containing species were broadly increased by palbociclib treatment and EMP2 gene disruption, the plurality of PUFA-containing species that were increased belonged to the plasmalogen-PE (PE P) class (Figure S7B). Furthermore, every PE P and PE lipid that was significantly altered contained a PUFA tail group (Figure S7B).

To analyze the dataset more broadly, we performed Bioinformatics Methodology for Pathway Analysis (BioPAN)^46^ on the lipidomics dataset. BioPAN revealed similarities between the pathways altered by palbociclib in Control cells, and pathways altered by EMP2^KO^ cells compared to Control (Figure S7C). Synthesis of PE phospholipids from PS lipids was enriched in both conditions, while catabolism of PE to PS was disenriched. Synthesis of PE P lipids was highly enriched (z-score > 4) in both ferroptosis-sensitizing conditions (Figure S7C). Next, we performed principal component analysis (PCA) and found that the Control + palbociclib and EMP2^KO^ + DMSO conditions clustered closely together (Figure 6G). Specifically, these conditions were primarily separated from the Control + DMSO samples in principle component 1 (Figure 6G). Further analysis of the PCA loadings plots revealed that compared to phosphatidylcholines, which were dispersed across principal component 1, PE and PE P lipids appeared to contribute to an increase in principal component 1 (Figure 6H).

We therefore more closely investigated the specific PE and PE P lipids altered by palbociclib treatment and EMP2^KO^ (Figures S7A and S7B). The only PE lipid that met the statistical criteria was PE 16:0_20:4 (Table S5). Of the seven PE P lipids significantly altered, a plurality (3 of 7) contained a 20:4 tail group: PE P-16:0/20:4, PE P-18:0_20:4, and PE P-22:0/20:4 (Table S5). All four 20:4-containing PE and PE P lipids were increased to the same extent in the Control + Palbociclib, EMP2^KO^ + DMSO, and EMP2^KO^ + palbociclib conditions compared to the Control + DMSO condition (Figure 6I and 6J). Thus, both palbociclib treatment and EMP2^KO^ enrich for pro-ferroptotic 20:4 tail groups in PE and PE P phospholipids, consistent with a lipid-dependent mechanism of ferroptosis sensitization.

### CDK4/6 inhibition and GPX4 inhibition enhance lipid peroxidation in vivo

Existing covalent GPX4 inhibitors such as RSL3 are unsuitable for use in vivo due to poor bioavailability. From the patent literature, we identified a novel orally bioavailable GPX4 inhibitor, compound 28 (Figure 7A).^47^ We confirmed that compound 28 potently induced cell death in cultured HT-1080^N^ cells and that this lethality was enhanced by palbociclib pretreatment (Figure 7B). Cell death induced by compound 28 was suppressed by co-treatment with ferrostatin-1, consistent with the induction of ferroptosis (Figure 7C).

**Figure 7.**
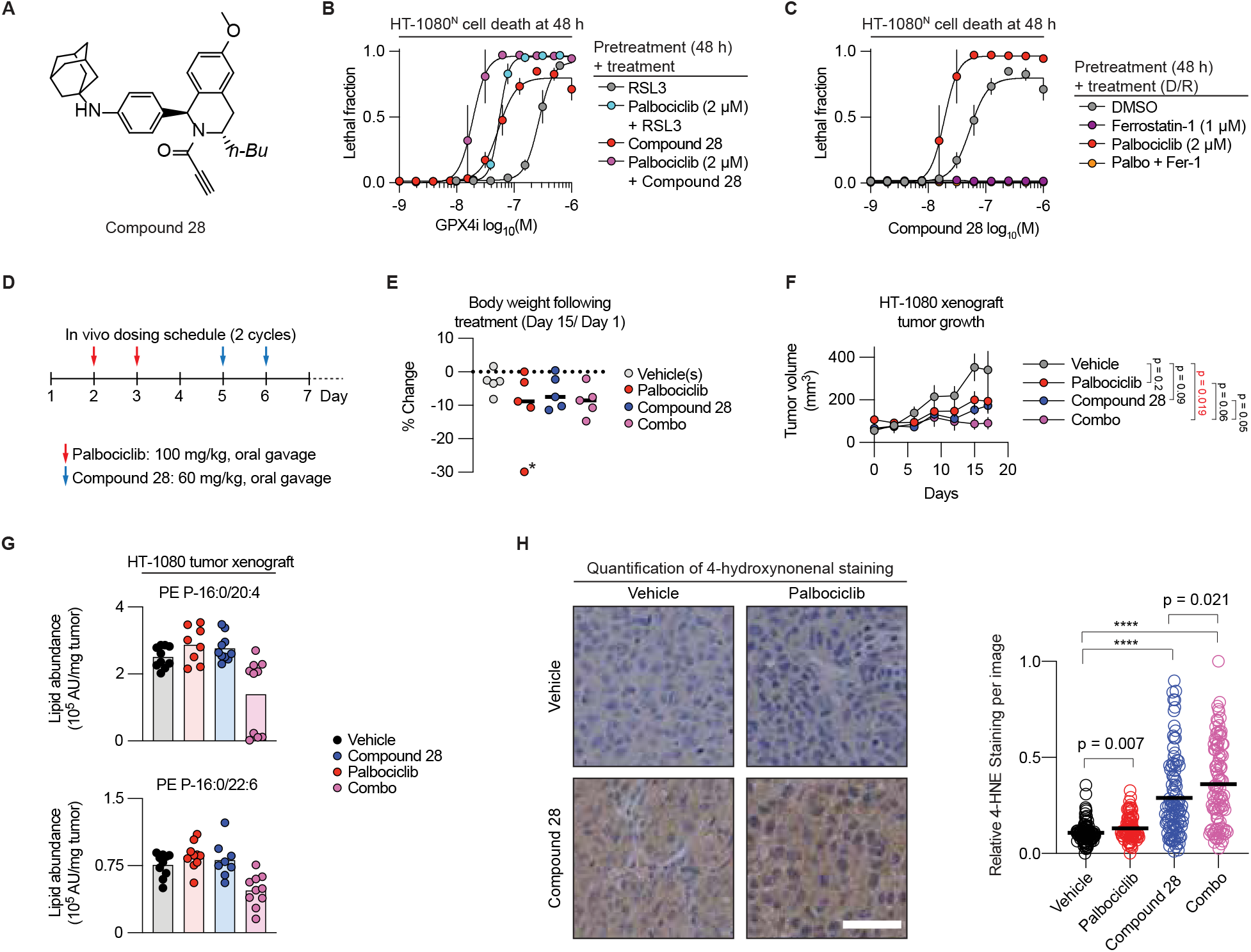
CDK inhibition modulates ferroptosis in vivo. (A) Structure of compound 28. (B, C) Cell death measured by imaging following pretreatment. Note that the compound 28 data in the DMSO and palbociclib pre-treatments conditions (without ferrostatin-1) in (C) are the same as shown in (B) as this was all part of one larger experiment. Data represent mean ± SD from three independent experiments. (D) A dosing schedule for the combination of palbociclib and compound 28. (E) Change in mouse body weight from the start to the end of the xenograft study. Body weight was measured every 3 d. Each dot represents data from one animal. Note: two xenograft tumors were implanted per animal, one on each flank. *One animal that lost body weight rapidly between days 12 and 15 was not included in the subsequent analysis. (F) Tumor xenograft size over time, determined by caliper measurements. Data are mean ± SEM, n = 8-10 tumors per condition. p-values indicate the results of 2-tailed t-tests of tumor volumes on day 17. (G) Lipid abundance measured by liquid chromatography coupled to mass spectrometry. Individual datapoints from 8 or 10 individual tumors. AU, arbitrary units. (H) Staining and quantification of the lipid peroxidation breakdown product 4-hydroxynonenal (4-HNE) in tumor xenografts. Scale bar = 50 µm. In the graph, each dot represents quantified intensity from one region of interest (ROI) from a xenograft tissue section. Ten ROIs/xenograft sample were acquired. Horizontal black bars indicate mean for each condition. ****2-tailed t-test, p < 10^-10^.

Both palbociclib and compound 28 independently have the potential to cause significant on-target toxicities in animals (e.g., kidney toxicity upon GPX4 disruption).^48^ We developed a weekly oral dosing schedule for palbociclib pre-treatment following by compound 28 treatment that was generally tolerated by mice over 17 days, albeit with a degree of weight loss (Figures 7D and 7E). Only mice treated with the combination of palbociclib and compound 28 (combo) had reduced tumor sizes at the experimental endpoint relative to untreated controls (Figure 7F). Consistent with this, tumors extracted from mice that received the combo treatment showed significantly lower levels of PE P-16:0/20:4 and PE P-16:0/22:6 (Figure 7G). Moreover, histologic analysis of the accumulation of 4-hydroxynonenal, a breakdown product of lipid peroxidation, was significantly higher in xenografts from animals that received both palbociclib and compound 28 than xenografts exposed to either compound alone (Figure 7H). These results suggest that a combination of CDK inhibition and GPX4i may induce more ferroptosis in tumor cells in vivo.

## DISCUSSION

Depending on how ferroptosis is triggered, different genes and mechanisms appear important for establishing sensitivity or resistance.^12^ We are only beginning to identify and characterize these trigger-specific regulatory mechanisms. Consistent with our own and other recent findings,^16–18^ we show here p53 stabilization suppresses ferroptosis induced by system x_c_^-^ blockade yet simultaneously enhances ferroptosis triggered by direct GPX4 inhibition. In addition, we demonstrate that inhibition of CDK4/6 activity is sufficient to selectively sensitize to GPX4i. Selective GPX4i sensitization requires ACSL4-dependent PUFA metabolism. This is consistent with findings that ACSL4 is more important for ferroptosis induced by direct GPX4 inhibition than by system x_c_^-^ inhibition.^12^ These results indicate that activation status of the CDK4/6 pathway may be one important determinant of sensitivity to distinct modes of ferroptosis induction.

Our findings are not entirely in agreement with other recent findings. For example, it has been reported that CDKN1A expression can suppress ferroptosis in some contexts.^18, 20^ A key difference between our results and other studies is the use of a lethal small molecule (named CETZOLE) that may act as both a system x_c_^-^ inhibitor and a covalent inhibitor of GPX4 and other proteins.^18^ As we find here, p53 and p21 expression can have opposite effects on ferroptosis depending on the induction mechanism (system x_c_^-^ inhibition or direct GPX4 inhibition) and this distinction may be important for understanding discrepancies between our results and previous findings. Additionally, we note that the time-lapse imaging approach used in our study may allow us to more carefully differentiate cell death from proliferative arrest compared to readouts of cell viability used in other studies.^21, 49^ Alternatively, differences in cell growth conditions or other biological factors could explain the differences between our results and published findings.

Many different lipids and lipid metabolic enzymes can contribute to ferroptosis in a context-dependent manner.^12^ An emerging concept is that ferroptosis sensitivity is not dictated by any one specific lipid species but rather by the overall abundance of PUFA-containing versus MUFA-containing phospholipids.^8, 12, 28, 38^ Consistent with a recent report,^35^ we find that overexpression of MBOAT1 can inhibit ferroptosis, likely by increasing the incorporation of MUFAs instead of PUFAs into membrane phospholipids, especially PEs.^7, 50^ However, these findings are correlative, and other lipid species not detected in our lipidomic analysis could explain the modulation of ferroptosis sensitivity. Interestingly, disruption of MBOAT1 alone was insufficient to sensitize to ferroptosis in HT-1080^N^ and other cell lines. This is likely explained by redundancy with a related enzyme, MBOAT2.^35^ It may be that the effect of MBOAT1 loss on lipid metabolism is most pronounced at the G1/S transition and therefore only detectable in cells that are effectively synchronized by cell cycle arrest. An interesting question for future exploration is the role of MBOAT1 in the cell cycle. Alterations of the lipidome are known to occur over the course of the cell cycle^51, 52^ and we speculate that changes in *MBOAT1* expression could link cell cycle phase to membrane lipid remodeling necessary for cell division.

Our results pinpoint *EMP2* as a cell cycle-regulated gene and important regulator of ferroptosis sensitivity. EMP2 is a lipid raft-associated cell surface protein, and in this capacity may help orchestrate lipid metabolism relevant to the induction of ferroptosis. Loss of *EMP2* expression increases the levels of PUFA-PL species that are linked to the induction of ferroptosis, providing a plausible mechanistic link between cell cycle arrest, *EMP2* downregulation, phospholipid remodeling and enhanced sensitivity to GPX4i. How EMP2 expression alters membrane phospholipid composition will require further elucidation but could involve interactions with plasma membrane-localized lipid remodeling enzymes or effects on lipid trafficking. Anti-EMP2 inhibitory antibodies have been shown to control tumor xenograft growth,^53, 54^ and it may be interesting in the future to examine combinations of these agents with GPX4i for cancer therapy.

We have explored the potential for combination treatments involving direct GPX4 inhibition in vivo. We have made use of a clinical-candidate GPX4 inhibitor (compound 28) reported in the patent literature. Compound 28 only had a statistically significant effect on HT-1080 xenografts tumor volume when combined with palbociclib administration. Moreover, palbociclib enhanced the levels of lipid peroxidation induced by compound 28. Another GPX4 inhibitor, compound 24, was also recently described.^27^ Compound 24 was tolerated by mice but had little effect alone on the growth of a diffuse large B cell lymphoma xenograft. It may be that GPX4 inhibition alone is not sufficient in many cases to affect tumor growth. Of note, compound 28 can be delivered via oral gavage, a potential advantage over compound 24. Toxicity is a concern with all GPX4 inhibitors^48^ and we did observe body weight loss in mice treated with compound 28. This toxicity will have to be carefully monitored in future studies. However, the existence of compound 28 opens a path to these and many other future studies.

### Limitations of the study

*MBOAT1* knockdown or gene-disruption is not sufficient to sensitize to GPX4 inhibition alone, and this may be explained by redundancy with *MBOAT2*.^35^ The correlation between *EMP2* gene expression and ferroptosis sensitivity are strong. However, we have been unable to identify a protocol that would allow us to detect EMP2 protein levels change in response upon cell cycle arrest in our cells of interest. The mechanistic link between *EMP2* expression and changes in lipid metabolism remains to be defined in detail. In vivo combination studies are challenging when using two drugs that each have potential dose-limiting toxicities. The compound 28 dosing strategy employed here may only be effective in this context and require further optimization for different cell models and drug combinations.

## SIGNIFICANCE

It may be possible to target the ferroptosis mechanism to treat cancer and other diseases. Emerging evidence indicates that system x_c_^-^ inhibition and direct inhibition of GPX4 do not induce ferroptosis in the same manner.^11^ Here, we show that stabilization of p53 and inhibition of CDK activity create unique cell states that exhibit distinct responses to cystine deprivation versus direct GPX4 inhibition. We show that cell cycle arrest is associated with transcriptional downregulation of several genes, including *MBOAT1* and *EMP2*, which can regulate the abundance of ferroptosis-associated lipids and thereby alter sensitivity to GPX4 inhibition. Using an orally bioavailable GPX4 inhibitor we also showed that combined CDK4/6 and GPX4 inhibition can enhance lipid peroxidation in tumors in vivo. These developments could help open new avenues to investigate ferroptosis in vivo.

## ACKNOWLEDGEMENTS

We thank P. Beltran and C. Stahlhut for providing compound 28 and advice, S. Terrel for assistance with lipidomics experiments, T. Hammond for creating lentiviral vectors, L. Pope and W. Lee for experimental assistance, E. Peterson for performing IHC, N. Terrell for assistance with computational analyses, J. Carette for reagents, J. Skotheim for advice, and L. Magtanong and members of the Dixon lab for comments on the manuscript. Certain constructs were obtained from Addgene. J.A.O. is a Chan Zuckerberg Biohub – San Francisco Investigator. This work was supported by funding from Ferro Inc. to S.J.D., awards from the National Institutes of Health to J.R. (1F31CA265146), Z.T.S., (R01CA262439), J.A.O. (R01GM112948) and S.J.D. (2R01GM122923), and an award from the American Cancer Society (RSG-21-017-01-CCG) to S.J.D.

## AUTHOR CONTRIBUTIONS

Conceptualization, J.R., M.L., J.H., A.T., Z.T.S., J.A.O., J.S., J.Z.L., S.J.D.; Methodology, J.R., V.L.L., A.H., M.L., J.H., A.T.; Investigation, J.R., N.K., V.L.L., A.H., M.L., J.H., A.T.; Writing, J.R., and S.J.D.; Funding Acquisition and Supervision, Z.T.S., J.A.O., J.S., J.Z.L., and S.J.D.

## STAR METHODS

### Chemicals and reagents

ML162 was synthesized by Acme (Palo Alto, CA). Erastin2 (Cat# 27087) and deferoxamine (Cat# 14595) were obtained from Cayman Chemicals (Ann Arbor, MI). Palbociclib (Cat# S1116), pimasertib (Cat# S1475), and 1*S*,3*R*-RSL3 (simply, RSL3) (Cat# S8155) were obtained from Selleck Chemicals (Houston, TX). Doxycycline (Dox) (Cat# D3447), ML210 (Cat# SML0521), ferrostatin-1 (Cat# SML0583) and propidium iodide (Cat# P4170) were obtained from Sigma-Aldrich (St. Louis, MO). Abemaciclib (Cat# HY-16297A) was obtained from MedChem Express (Princeton, NJ). Bortezomib (Cat# NC0587961), Q-VD-OPh (Cat# OPH00101M), etoposide (Cat# 12-261-00), DAPI (Cat# D1306) and C11 BODIPY 581/591 were from Thermo Fisher Scientific (Waltham, MA). Compound 28 was synthesized as described^47^ and provided as a gift by P. Beltran (Ferro Inc). Dox was dissolved in deionized water and stored at −20°C. C11 BODIPY 581/591 was dissolved into methanol and stored at −20°C. All other compounds were dissolved in dimethyl sulfoxide (DMSO) and then stored at −20°C. The compound library (Cat# L1700) was obtained from Selleck Chemicals^8^.

### Cell culture

HT-1080^N^, Caki-1^N^, H1299^N^, T98G^N^, and A375^N^ cells were described previously.^16, 21^ HT-1080^N^ Control and gene-disrupted p53^KO^ cells were described previously.^16^ HT-1080^N^ rtTA-CDKN1A cell lines were described.^17^ HT-1080^N^ Control and ACSL4^KO1/2^ cell lines were described previously.^12^ H460 Control and gene disrupted FSP1^KO^ cell lines were described previously.^55^ A HAP1 *MBOAT1* gene-disrupted cell line and a paired wild type control cell line were purchased from Horizon Discovery (Cambridge, UK). 786-O and MDA-MB-231 cells were obtained from ATCC (Manassas, VA). The generation of HT-1080^N^ HA-CCND1^WT^ and HT-1080^N^ HA-CCND1^T286A^, HT-1080^N^ rtTA-MBOAT1^WT^ and HT-1080^N^ rtTA-MBOAT1^H381A^, and HT-1080^N^ Control and EMP2^KO^ cells are described below. All HT-1080-based cell lines were cultured in DMEM Hi-glucose medium (Cat# MT-10-013-CV, Corning Life Science, Corning, NY) supplemented with 10% fetal bovine serum (FBS, Cat# 26140-079, Gibco/Thermo Fisher Scientific), 1% non-essential amino acids (NEAAs) (Cat# 11140-050, Life Technologies, Carlsbad, CA), and penicillin + streptomycin (P/S) (5000 U/mL) at 1x concentration (Cat# 15070-063, Life Technologies). H460, HAP1^N^, T98G^N^, H1299^N^, 786-0 and MDA-MB-231 cells were grown in DMEM Hi-glucose medium supplemented with 10% FBS and 1x P/S. A375^N^ and Caki-1^N^ cells were cultured in McCoy’s 5A media (Cat# MT-10-050-CV, Corning Life Science) supplemented with 10% FBS and 1x P/S. All cells were grown in humidified incubators (Thermo Fisher Scientific) at 37°C with 5% CO2.

### Cell death measurement

Cells were seeded to obtain 70-80% confluency at the time of lethal stimulus addition. All HT-1080-based cell lines and Caki-1^N^ cells were seeded at 1,500 cells/well in DMSO treatment and at 3,000 cells/well in 96-well plates for treatment with nutlin-3, palbociclib, pimasertib, or abemaciclib. A375^N^ cells were seeded at 3,000 cells/well for DMSO conditions and at 6,000 cells/well in 96-well plates for nutlin-3 and palbociclib treatment conditions. HAP1^N^ parental and MBOAT1^KO^ cells were seeded at 2,500 cells/well for DMSO conditions and 5,000 cells/well in 96-well plates for palbociclib treatment. HT-1080^N^ rtTA-CDKN1A cells were seeded at 1,500 cells/well for DMSO treatment, and 3,000 cells/well in 96-well plates for palbociclib, Dox, and Dox + palbociclib treatment. HT-1080^N^ rtTA-MBOAT1^WT^ and HT-1080^N^ rtTA-MBOAT1^H381A^ cells were seeded at 1,500 cells/well for DMSO and Dox treatment, and 3,000 cells/well in 96-well plates for palbociclib and Dox + palbociclib treatment. The next day, cells were treated with the indicated pretreatment. For these cell lines, the concentrations of treatment compounds were as follows: palbociclib (2 μM), nutlin-3 (10 μM), abemaciclib (1 μM), pimasertib (0.5 μM), Dox (0.1 μg/mL). 48 h after pretreatment was initiated, cells were exposed to a lethal stimulus (e.g., RSL3). Additionally, SYTOX Green (20 nM, Cat# S7020, Life Technologies/Thermo Fischer Scientific) was added to the growth medium to track cell death. Images were acquired using either an IncuCyte ZOOM imaging system (Model 4459, Essen BioSciences, Ann Arbor, USA) or an IncuCyte S3 live cell analysis instrument (Sartorius, Göttingen, Germany) housed within a tissue culture incubator maintained at 37°C with 5% CO2. In each well, phase contrast, red (excitation: 585 ± 20, emission: 665 ± 40, acquisition time: 800 ms), and green (excitation: 460 ± 20, emission: 524 ± 20, acquisition time: 400 ms) images were obtained at 10x magnification. The images were analyzed using either the ZOOM software package (V2016A/B) or the S3 software package. Lethal fraction was calculated as described.^56^

Cell death studies of 786-O and MDA-MB-231 cells were conducted as follows. Cells were seeded in 10 cm plates at the density of 800,000 cells/plate and treated with vehicle control or palbociclib (1 µM) for 48 h. After pretreatment, cells were trypsinized, seeded into 12-well plates (80,000 cells per well), and treated with ferroptosis-inducing compounds. At the end of treatment, the culture medium was collected. Cells were then washed with 300 μL PBS and the PBS was combined with culture medium. Cells were then trypsinized using 100 μL trypsin and neutralized with the collected culture medium. After centrifugation at 500 x *g* for 5 min, cells were resuspended in 180 μL PBS containing 2 μg/mL propidium iodide (PI) and PI-positive dead cells were detected by flow cytometry using a BD fortessa X-20 cytometer. Percent cell death was determined as the percentage of PI-positive cells in the sampled population.

### Small molecule modulation screen

HT-1080^N^ cells were seeded into 384-well plates at a density of 1,000 cells per well. The next day, either DMSO or nutlin-3 (final concentration: 10 µM) was added to the growth media. 48 h after the addition of DMSO or nutlin-3, a library of 261 bioactive compounds was added to each plate (final concentration: 5 µM) using a Versette automated liquid handler as described.^8^ Images were acquired every 4 h using the IncuCyte ZOOM as described above. Lethal fraction scores for each time point were calculated as above, using custom Python code. Area-under-curve (AUC) analysis was performed in Python measuring the AUC of the lethal fraction versus time. AUC values were normalized to the total time of the experiment (nAUC). For each bioactive compound, the nAUC of each test compound alone was compared to the nAUC of nutlin-3 alone, as well as the nutlin-3 + compound condition. Compound-compound interactions were assessed using the Bliss Independence model, yielding an interaction score for each compound. Z-scores were computed across the set of all compound interaction scores and plotted.

### Cloning

The coding sequence (CDS) of *MBOAT1* (NM_001080480.3) was obtained from NCBI. A gene block was designed such that the CDS was flanked on either side by overhangs useful for subsequent cloning. At the 5’ end, a Kozak consensus sequence was inserted immediately prior to a sequence encoding a 1x FLAG tag and a 15-nucleotide spacer (5’-ACTAGTCCAGTGTGGTGGAATTCTGCAGATACCATGGATTACAAGGATGACGACGATAAGGCCCGGGCGGATCCC-3’). The 3’ overhang sequence was as follows (5’-ATCCAGCACAGTGGCGGCCGCTCGACAATC-3’). The final sequence (hereafter denoted “MBOAT1 insert”) was confirmed by Sanger sequencing. The destination plasmid pLenti TRE3G Dest (pSD292) was a generous gift of Jan Carette (Stanford School of Medicine). pSD292 was linearized by digestion with *EcoRV* (Cat# R3195S, New England Biolabs, Ipswich, MA) as per manufacturer protocol. The MBOAT1 insert was cloned into pSD292 through a Gibson assembly protocol wherein 100 ng of linearized pSD292 (19.9 fmol) and a 3x molar ratio of insert (59.7 fmol) were combined with the Gibson assembly mix (Cat# E5510S, New England Biolabs) per manufacturer instructions. The new plasmid, termed “pLenti TRE3G-MBOAT1”, was transformed into NEB 5-alpha competent *E. coli* provided with the Gibson assembly kit (Cat# E5510S, New England Biolabs) per manufacturer protocol. pLenti TRE3G-MBOAT1^H381A^ was generated by using the Q5 mutagenesis kit (Cat# E0554S, New England Biolabs). In brief, forward (5’-TGCTTTGTGGgcTGGTGTCTAC-3’) and reverse (5’-GACAGGATGAAGGTTAGC-3’) primers were added to the master mix at 10 µM each. The annealing temperature was 56°C. All other steps were performed per manufacturer instructions. pLenti TRE3G-MBOAT1^H381A^ was transformed into NEB 5-alpha competent *E. coli* as above.

### Lentivirus production

293T cells were seeded in 6-well plates (Cat# 3516, Corning) at a density of 500,000 cells/well. The next day, medium was removed and replaced with DMEM lacking P/S. Cells were transfected using 1,000 ng of plasmid DNA for pLenti TRE3G-MBOAT1 or pLenti TRE3G-MBOAT1^H381A^, together with the lentiviral packaging plasmids pMD2.G (250 ng, Cat# 12259, Addgene, Watertown, MA) and psPax2 (750 ng, Cat# 12260, Addgene) using Lipofectamine LTX transfection reagents (Cat# 15338-100, Life Technologies) diluted in OptiMEM media (Cat# 31985070, Thermo Fisher Scientific) to a final volume of 250 µL. The transfection mixture was incubated for 15 min and then added dropwise to the cells. The cells were then incubated overnight. The next day, virus-containing medium was collected from both plates and replaced with 2 mL of fresh medium. 8 h later, the medium was collected and replaced with 2 mL fresh medium. The following morning, the medium was collected a final time (for a total of 6 mL virus-containing medium per virus). The viral media was filtered using a 0.45 μm filter (Cat# SLHV033RS, Millex/Sigma Aldrich, St. Louis, MO), aliquoted, and frozen at −80°C until use.

### Generation of CCND1-overexpressing cells

Lentiviruses that express the vector for HA-CCND1^WT/T286A^ were described.^57^ HT-1080^N^ cells were seeded in a 6-well dish such that each well contained 50,000 cells (day 0). On day 1, cells were treated with lentivirus-containing growth medium containing 8 µg/mL polybrene at a viral MOI of 1. On day 2, medium was removed, and puromycin was added in fresh growth medium at a final concentration of 1 μg/mL. On day 4, medium was replaced with fresh medium lacking puromycin, and the surviving cells were used in subsequent experiments.

### Generation of MBOAT1-overexpressing cell lines

HT-1080^N^ rtTA cells,^17^ were seeded at 3,000 cells/well in 96-well plates (Cat# 781660, BrandTech, Essex, CT). The next day, cells were infected with a serial dilution of either MBOAT1^WT^ or MBOAT1^H381A^ lentiviral particles in growth media containing 8 µg/mL polybrene (Cat# H9268-5G, Sigma Aldrich). 24 h later, viral media were removed and replaced with media containing 1 μg/mL puromycin (Cat# A11138-03, Thermo Fisher Scientific). Cells infected with 10% virus-containing media completely survived puromycin selection. Therefore, HT-1080^N^ rtTA cells were infected using 10%/90% virus-containing medium/regular growth medium containing 8 µg/mL polybrene. 24 h later, viral medium was removed, 1 μg/mL puromycin was added for 48 h to eliminate uninfected cells. The cell lines were expanded into cultures referred to as HT-1080^N^ rtTA-MBOAT1^WT^ and HT-1080^N^ rtTA-MBOAT1^H381A^.

### Generation of EMP2^KO^ cells

pSpCas9(BB)-2A-GFP plasmid was obtained from Addgene (Cat# 48138). sgRNA primers for EMP2 were synthesized as follows. The forward primer for the sgRNA was 5’-GAAACTTACATTGTCGACGG and the reverse primer was 5’-CCGTCGACAATGTAAGTTTC. 5’ and 3’ adaptors were added to the primers to match the *Bsp*I cleavage sites of pSpCas9(BB)-2A-GFP such that the final primers were F: 5’ CACCGGAAACTTACATTGTCGACGG and R: 5’-AAACCCGTCGACAATGTAAGTTTCC. The primers were diluted to a final concentration of 10 µM in 10 µL of H_2_O and then heated to 95°C, cooled to 25°C at a rate of 2.5°C/min. The duplexed oligo was diluted 1:200 in H_2_O and then ligated into the backbone through the following reaction. 100 ng backbone, 2 µL of the diluted oligo duplex, 2 µL of 10x FastDigest buffer (Thermo Fisher Scientific), 1 µL 10 mM DTT, 1 µL 10 mM ATP, 1 µL FastDigest *Bpi*I (Cat# D1014, Thermo Fisher Scientific), 0.5 µL T4 DNA ligase (Cat# EL0014, Thermo Fisher Scientific), and up to 20 µL of H_2_O were combined. The mixture was incubated for six cycles of 37°C, 5 min then 21°C, 5 min. The product was transformed into competent DH5α cells as above, and purified plasmid was collected from the cells using the Qiagen Plasmid Plus Midi Kit (Cat# 12943) per manufacturer protocol.

HT-1080^N^ cells were transfected with 1 μg of plasmid using Lipofectamine LTX reagent as described above. Briefly, 50,000 cells were seeded into a well of a 6-well plate. Cells were transfected, and then transfection media was removed the following morning. 24 h later, cells were trypsinized and sorted at the Stanford FACS facility. GFP+ cells were sorted into a 96 well plate containing 200 μL of media that contained 30% FBS such that only one cell was deposited into each well. Homogenous populations were allowed to grow for three weeks, and then each clone was expanded and the *EMP2* locus was sequenced. A wild-type control (“Control”) was isolated as well as an independent clone contained a homozygous 1-bp deletion resulting in a premature stop codon seven amino acids downstream that was named EMP2^KO^.

### Cell cycle profiling

All cells were seeded to obtain 70-80% confluency in 6-well plates at time of harvest. Cells treated with DMSO were seeded at 30,000 cells per well. Cells treated with nutlin-3, palbociclib, pimasertib, or abemaciclib were seeded at 70,000 cells per well. HT-1080^N^ rtTA-CDKN1A cells treated with Dox or Dox + palbociclib were seeded at 70,000 cells per well, whereas HT-1080^N^ rtTA-MBOAT1^WT^ and rtTA-MBOAT1^H381A^ cells treated with Dox were seeded at 30,000 cells per well. HT-1080^N^ rtTA-MBOAT1^WT^ and rtTA-MBOAT1^H381A^ cells treated with Dox + palbociclib were seeded at 70,000 cells per well. HAP1^N^ MBOAT1 WT and KO cells were seeded at 50,000 cells per well in DMSO conditions and 100,000 cells per well for palbociclib treatment. The next day, cells were treated with the indicated conditions for 48 h prior to analysis. For all cell lines, the concentrations of treatment compounds were as follows: palbociclib (2 μM), nutlin-3 (10 μM), abemaciclib (1 μM), pimasertib (0.5 μM), Dox (0.1 μg/mL). Cells were then trypsinized using 0.25% trypsin-EDTA (Cat# 25200114, Thermo Fisher Scientific) and washed 2x with 1.5 mL cold HBSS (Cat# 14025-134, Life Technologies). Cells were fixed in 70% ethanol and stored at −20°C for up to two weeks. Immediately prior to analysis, cells were brought to room temperature, washed 2x with 1.5 mL cold HBSS, and stained with solution containing RNase A solution (1:100 v/v, Cat# 19101, Qiagen, Hilden, Germany) at and propidium iodide (PI) solution (1:20 v/v, Cat# P3566, Thermo Fisher Scientific) for 30 min at 37°C. After staining, cells were washed 1x with 1 mL of HBSS, resuspended in 500 µL HBSS, and strained through a filter cap FACS tube (Cat# 352235, Corning). Cell cycle status was quantified on an Attune NxT Flow Cytometer. Data was acquired using standard laser excitation (488 nm) and emission (filter 590/40 nm) settings. Cells were assigned to G1, S, and G2 phase based on PI fluorescence histograms analyzed with the FlowJo v10.8.1 cell cycle analysis tool and Dean-Jett-Fox model as described (https://docs.flowjo.com/flowjo/experiment-based-platforms/cell-cycle-univariate/).

### Western blotting

All cells were seeded in 6-well plates to obtain 70-80% confluence at time of harvest. Cells treated with DMSO were seeded at 30,000 cells per well. Cells treated with nutlin-3, palbociclib, pimasertib, or abemaciclib were seeded at 70,000 cells per well. HT-1080^N^ rtTA-CDKN1A cells treated with Dox or Dox + palbociclib were seeded at 70,000 cells per well, whereas HT-1080^N^ rtTA-MBOAT1^WT^ and rtTA-MBOAT1^H381A^ cells treated with Dox were seeded at 30,000 cells per well. HT-1080^N^ rtTA-MBOAT1^WT^ and rtTA-MBOAT1^H381A^ cells treated with Dox + palbociclib were seeded at 70,000 cells per well. The next day, cells were treated with various drugs/conditions. For all cell lines, the concentrations of treatment compounds were as follows: palbociclib (2 μM), nutlin-3 (10 μM), abemaciclib (1 μM), pimasertib (0.5 μM), Dox (0.1 μg/mL). After 48 h, cells were washed 2x with 1.5 mL of cold HBSS and then scraped with a cell scraper (Cat# 07-200-364, Fisher Scientific). Cells were then pelleted and lysed in RIPA lysis buffer + 0.1% SDS with 1:200 protease inhibitor cocktail P8340 (Cat# P8340, Sigma-Aldrich). Harvested materials were incubated on ice for at least 30 min to ensure full lysis and then sonicated 10x at 20% amplitude using a Fisher Scientific Model 120 sonic dismembrator (Thermo Fisher Scientific). The lysates were centrifuged for 15 min at 18,213 x *g* at 4°C and supernatants were transferred to a new Eppendorf tube. Then, protein levels were quantified using the bicinchoninic acid (BCA) reagent (Cat# 23228, Thermo Fisher Scientific) per the manufacturer protocol in 96-well plate format. For the BCA assay, absorbance at 562 nm was measured using a Synergy Neo2 multimode plate reader (BioTek Instruments, Winooski, VT). Samples to be analyzed were prepared using 4x Laemmli Sample Buffer (Cat# 1610747, Bio-Rad, Hercules, CA) and run on 4-15% Mini-PROTEAN TGX precast protein gels (Cat# 4560184, Bio-Rad) in Tris-glycine running buffer (Cat# 1610772, Bio-Rad) at 100V. Proteins were transferred to nitrocellulose membranes using the iBlot2 system (Life Technologies). Blots were blocked for 1 h at room temperature in Intercept blocking buffer (Cat# 927-60001, LI-COR Biosciences, Lincoln, NE) incubated in primary antibody solution overnight at 4°C. Antibodies used were anti-ACSL4 (1:300, Cat# 22401-1-AP, Proteintech, Rosemont, IL), anti-Actin (1:4,000, C4, Cat# sc-47778, Santa Cruz Biotechnology, Dallas, TX), anti-Tubulin (1:5,000, DM1A, Cat# MS581P1, Thermo Fisher Scientific), anti-p21 (1:500, 12D1, Cat# #2947, Cell Signaling Technology, Danvers, MA), anti-Rb (1:1,000, G3-245, Cat# 554136, BD Biosciences, San Jose, CA), anti-phospho-Rb Ser807/811 (1:500, D2OB12, Cat #8516, Cell Signaling Technologies), anti-p53 (1:1,000, DO-1, Cat# sc-126, Santa Cruz Biotechnology), anti-GPX4 (1:500, Cat# ab41787, Abcam) anti-GPX4 (1:1,000, EPNCIR144, Cat# 125066, Abcam), anti-eIF4E (1:1000, P-2, Cat# sc-9976, Santa Cruz Biotechnology), anti-xCT (1:500, D2M7A, Cat# 12691, Cell Signaling Technology), anti-FLAG (1:1,000, Cat# ab1257, Abcam), anti-HA tag (1:1000, C29F4, Cat# 3724, Cell Signaling Technology), and anti-FSP1 (1:1000, Cat# 20886-1-AP, Proteintech). All antibodies were diluted into 5 mL of Intercept blocking buffer. The next day, membranes were washed 3x for 10 min with Tris-buffered saline (TBS, Cat# 0788, ISC BioExpress, Kaysville, UT) containing 0.1% Tween 20 (TBST) at room temperature. Membranes were incubated in secondary antibody solution at room temperature for 45 min at a 1:15,000 dilution in 1:1 Intercept blocking buffer:TBST. Antibodies used were 680LT Donkey-anti-mouse (Cat# 926-68022, LI-COR), 680LT Donkey-anti-rabbit (Cat# 926-68023, LI-COR), 800CW Donkey-anti-mouse (Cat# 926-32212, LI-COR), or 800CW Donkey-anti-rabbit (Cat# 926-32213, LI-COR). After secondary antibody incubation, blots were washed 3x for 10 min with TBST and then imaged on a LI-COR Odyssey CLx imager at 169 μm resolution, high quality setting, and 0.0 mm focus offset.

### RT-qPCR

Cells were seeded in a 6-well plate at densities such that at the time of cell harvest, plates were 70-80% confluent. Cells treated with DMSO were seeded at 30,000 cells per well. Cells treated with nutlin-3, palbociclib, pimasertib, or abemaciclib were seeded at 70,000 cells per well. HT-1080^N^ rtTA-CDKN1A cells treated with dox or dox + palbociclib were seeded at 70,000 cells per well, whereas HT-1080^N^ rtTA-MBOAT1^WT^ and rtTA-MBOAT1^H381A^ cells treated with dox were seeded at 30,000 cells per well. HT-1080^N^ rtTA-MBOAT1^WT^ and rtTA-MBOAT1^H381A^ cells treated with dox + palbociclib were seeded at 70,000 cells per well. The next day, cells were treated with the indicated conditions for 48 h prior to analysis. For all cell lines, the concentrations of treatment compounds were as follows: palbociclib (2 μM), nutlin-3 (10 μM), abemaciclib (1 μM), pimasertib (0.5 μM), dox (0.1 μg/mL). Next, cells were washed twice in warm PBS (0.5 mL) and then scraped into 0.5 mL PBS. Cells were lysed using a Qiashredder column (Cat# 79654, Qiagen). RNA was isolated using the RNeasy Plus RNA Extraction Kit (Cat# 74134, Qiagen) per manufacturer protocol and stored at −80°C. cDNA was generated with the TaqMan Reverse Transcriptase Kit (Cat# N8080234, Life Technologies) using 100 ng of RNA template. Quantitative PCR reactions were prepared with SYBR Green Master Mix (Cat# 4367659, Life Technologies) and run on an Applied Biosystems QuantStudio 3 real-time PCR machine (Thermo Fisher Scientific). Relative transcript levels were determined using the ΔΔCT method by normalizing the expression of the gene of interest to the expression levels of either *ACTB* or *GADPH*. qPCR primer sequences are provided in Table S4.

### Glutathione measurements

Total glutathione was assayed as described.^4^ 30,000 (DMSO) or 70,000 (nutlin-3 or palbociclib) HT-1080^N^ cells were seeded in 6-well plates. The next day, cells were treated with DMSO, nutlin-3 (10 µM), or palbociclib (2 µM) and then returned to the incubator for 48 h. 40 h later, a separate group of cells were treated with erastin2 (2 µM) for 8 h such that each condition ended its treatment time simultaneously. Following this treatment, cells were washed once with HBSS and scraped into MES buffer containing 1 mM EDTA. Samples were sonicated ten times with one second pulses at maximum amplitude on a Fisher Scientific Model 120 Sonic Dismembrator (Thermo Fisher Scientific) at 50% total amplitude. Lysates were centrifuged at >15,000 x *g* for 15 min at 4°C. Then samples were incubated for 5 min at room temperature in an equal volume of 12.5 M metaphosphoric acid (Cat# AC21922-1000, Thermo Fisher Scientific). Lysates were then centrifuged at >20,000 x *g* for 3 min at room temperature and the resultant supernatants were collected and stored at −20°C. Total glutathione was measured using the Cayman Glutathione Assay kit (Cat# 703002, Cayman Chemical) per the manufacturer’s instructions. After 30 min incubation, absorbance was measured using a Synergy Neo2 plate reader (BioTek) at 402 nm.

### C11 BODIPY 581/591 analysis

Confocal imaging was performed as described.^8^ Briefly, HT-1080 cells were seeded in 6-well tissue culture plates containing glass coverslips. 4 x 10^4^ cells were seeded for the DMSO condition and 8 x 10^4^ cells were seeded for the palbociclib condition. The next day, cells were treated with DMSO or palbociclib (2 μM) and returned to the incubator for 48 h. Then, DMSO or RSL3 (5 μM) was added for 1 h. Cells were then washed with pre-warmed HBSS. C11 BODIPY 581/591 (C11, 5 μM final concentration) and DAPI at 0.1 μg/mL final concentration were diluted in HBSS. 300 µL of the C11/DAPI/HBSS solution was added to cells for 10 min at 37°C. This solution was then aspirated and 1 mL of fresh HBSS was added to cells. Coverslips were mounted onto glass microscopy slides, which were imaged on a Zeiss Observer Z1 confocal microscope with a confocal spinning-disk head (Yokogawa, Tokyo, Japan), PlanApoChromat 63×/1.4 NA oil immersion objective, and a Cascade II:512 electron-multiplying (EM) CCD camera (Photometrics, Tucson, AZ). Three biological replicates were imaged per condition, with ten different regions of interest (ROI) captured per slide. Representative images are shown. Images were processed in ImageJ. Brightness for all images was auto adjusted based on images with the brightest signal.

For HT-1080^N^ rtTA-CDKN1A cells, C11 oxidation was detected using a BioTek Lionheart microscope. Here, cells were seeded in 96-well plates at 1,500 cells/well (for DMSO treatment) or 3,000 cells/well (for Dox, palbociclib, or Dox + palbociclib treatment). Two wells were seeded per condition. The next day, media were removed, and 200 μL of fresh growth media containing DMSO, Dox (0.1 μg/mL), palbociclib (2 µM), or Dox + palbociclib were added to the wells. 48 h later, media were removed and fresh media containing either DMSO or RSL3 (5 µM) were added to the cells such that each pretreatment condition (DMSO, Dox, palbociclib, Dox + palbociclib) had one well with DMSO and one well with RSL3. One hour later, media were removed from the wells, and each well was washed with 100 μL of warm sterile PBS. C11 (5 µM final concentration) and Hoechst (1 μg/mL final concentration) (Cat# H1399, Fisher Scientific) were diluted in warm PBS. 50 µL of the solution was added to each well for 10 min at 37°C. Then, the solution was removed and 100 μL PBS added to each well. The plate was immediately imaged on a BioTek Lionheart imager. The automated imager acquired one image per well using a 20x objective with a correction collar of 1.25 mm. DAPI (excitation: 377 nm, emission: 447), Texas Red (excitation: 586 nm, emission: 647), and GFP (excitation: 469 nm, emission: 525) filter cubes were used. Images were quantified using Gen5 v3.10 software (BioTek) to determine oxidized/non-oxidized (ox/non-ox) C11 ratio per cell. Ox/non-ox ratio is reported per each cell analyzed, summed across three replicates. More than 100 cells were quantified per condition.

For 786-O and MDA-MB-231 cells C11 oxidation was examined as follows. Cells were seeded in 10 cm plates at the density of 800,000 cells/plate and treated with vehicle control or palbociclib (1 µM) for 48 h. After pretreatment, cells were trypsinized, seeded into 12-well plates (80,000 cells per well), and treated with ferroptosis-inducing compounds ± inhibitors. At the end of treatment, the culture medium was collected. Cells were then washed with 300 μL PBS and the PBS was combined with culture medium. Cells were then trypsinized using 100 μL trypsin and neutralized with the collected culture medium. After centrifugation at 500 x *g* for 5 min, cells were resuspended in 300 μL PBS containing C11 (1 µM) for 30 min at 37°C. Then cells were resuspended in 180 μL PBS and detected by flow cytometry using BD fortessa X-20 cytometer. Live single cells were gated and the geometric means of FITC (green) and PE (red) intensity were acquired using FlowJo software. The green-to-red ratio was then calculated to indicate lipid peroxidation.

### RNA-sequencing

HT-1080^N^ cells were seeded in 6-well plates at 30,000 cells (DMSO) or 70,000 cells (palbociclib or pimasertib treatment). The next day, growth media was removed and replaced with 2 mL media containing DMSO, palbociclib (2 µM), or pimasertib (0.5 μM). After 48 h, RNA was extracted as described above. RNA quality was measured at the Stanford Protein and Nucleic Acid Facility using the Eukaryote Total RNA nanochip (Agilent Technologies). Library generation, read sequencing, and data cleanup were performed by Novogene.^49^ Briefly, an Illumina HiSeq 4000 platform was used to sequence bases, and reads were excluded if indeterminate bases consisted of >10% of the read or >50% of the bases had a quality score of less than five. The raw image files from the high-throughput sequencing were transformed into sequenced reads using CASAVA base recognition. Reads were then mapped to the reference genome (hg19) using the STAR aligner. Expression of each mRNA was calculated and reported as fragments per kilobase of transcript sequence per million base pairs sequenced (FPKM). The change in FPKM was compared between groups (e.g., DMSO versus palbociclib) and differential expression was analyzed using the DESeq2 R package. FDR correction was performed using the Benjamini-Hochberg method. Differentially expressed genes that met an FDR cutoff threshold of q < 0.05 were reported.

### siRNA transfection

siRNAs were purchased from Integrated DNA Technologies (IDT, Newark, NJ). The siRNA used were hs.Ri.MBOAT1.13.3, hs.Ri.EMP2.13.2 and hs.Ri.EMP2.13.3. AllStars Negative Control siRNA (siNTC, Cat# 1027280, Qiagen) was used as a validated non-targeting siRNA control for all experiments. HT-1080^N^ and Caki-1^N^ cells were seeded at 25,000 cells per well (in 12-well plates) or 75,000 cells per well (in 6-well plates). T98G^N^ and A375^N^ cells were seeded at 75,000 cells per well in 6-well plates, treated with hs.Ri.MBOAT1.13.3 for 48 h, and then trypsinized and seeded into 96-well plates at a density of 6,000 cells per well. The following day, the indicated doses of RSL3 were added to each well. For siNTC, siMBOAT1, and siEMP2, 10 pmol of siRNA per 250 μL of transfection reagent was used. siRNA was combined with 2.5 μL of Lipofectamine RNAiMAX transfection reagent (Cat# 13778075, Life Technologies) and 250 μL of OptiMEM reduced serum growth medium (Cat# 31985070, Thermo Fisher Scientific) in a new 12-well plate. For 6-well plates, the same ratios were used with double the amount of each reagent. The transfection mixture was incubated in the well for at least 15 min, and then cell suspensions were carefully added to each well. For relevant experiments/conditions, palbociclib was added at time of seeding at 2 μM. 48 h later, growth medium was removed and replaced with medium containing SYTOX Green (20 nM) and the indicated concentration of RSL3 and/or ferrostatin-1. For RNA measurements, cells were scraped 48 h after transfection and analyzed. (See “RT-qPCR” for relevant details.)

### Untargeted lipidomic analysis

HT-1080^N^ rtTA-MBOAT1^WT^ and HT-1080^N^ rtTA-MBOAT1^H381A^ cells were seeded in 10 cm plates. For both cell lines, 300,000 cells/plate were used for the DMSO and Dox treatment conditions, and 600,000 cells/plate were used for palbociclib or Dox + palbociclib treatment conditions. HT-1080^N^ ACSL4 WT, KO1, and KO2 cells were seeded in 10 cm plates. For all cell lines, 300,000 cells/plate were used for the DMSO condition, and 600,000 cells/plate were used for the palbociclib condition. One day after seeding, cell culture media were changed to fresh media containing DMSO, Dox (0.1 µg/mL), palbociclib (2 µM), or Dox + palbociclib. 48 h later, cells were washed 2x with 5 mL of warm PBS and then scraped into PBS using Corning cell lifters (Cat# 07-200-364, Fisher Scientific). 100 µL of cell extract was removed for protein quantification using the BCA assay (Cat# 23252, Thermo Fisher Scientific). Lipids were extracted from the cells using a modified Folch extraction, Final: 2:1:1 CHCl3:MeOH:PBS. Chloroform (Cat# 366927) and methanol (Cat# 34860-4L-R) were obtained from Sigma-Aldrich. All reagents used were HPLC grade. Lipid extracts were shaken vigorously for 10 s, and then centrifuged at 350 x *g* for 5 min at 4°C. The organic (lower) phase was removed into a fresh glass vial and then dried under N2. Lipids were stored at −80°C until run on the mass spectrometer.

On the morning of the analysis by mass spectrometry, lipids were resuspended in 100 µL of 9:1 MeOH:toluene. HPLC grade toluene was obtained from Sigma-Aldrich (Cat# 34866). The samples were then run as previously described.^58^ Briefly, buffer A was 60:40 acetonitrile (ACN):H2O, while buffer B was 90:10 isopropanol (IPA):ACN. In positive mode the additives were 0.1% formic acid in buffer A and 10 mM ammonium formate and 0.1% formic acid in buffer B. In negative mode, the additive was 10 mM ammonium acetate. Each run was for 15 min with the following conditions: 0 min 85% (A); 0-2 min 70% (A); 2-2.5 min 52% (A); 2.5-11 min 18% (A); 11-11.5 min 1% (A); 11.5-12 min 1% (A); 12-12.1 min 85% (A). For MBOAT1 overexpression experiments, an Acquity UPLC CSH C18 column (100 x 2.1 mm; 1.7 µm, Cat# 186005297) maintained at 65°C was used in for high-performance liquid chromatography (HPLC) analysis. For HT-1080^N^ Control and *ACSL4* gene-disrupted cells, an Agilent ZORBAX RR Eclipse Plus C18 column (95Å, 4.6 x 100 mm, 3.5 µm, part number 959961-902) maintained at 65°C was used in the HPLC phase. For HPLC runs on this column, an additional five minutes of equilibration was added (15-20 min) at 85% (A). Samples were maintained at 4°C in the autosampler for the duration of the experiment.

An Agilent (Santa Clara, CA) 6545 quadrupole time of flight (qTOF) mass spectrometer was used to acquire data in negative electrospray ionization mode with the parameters: mass range *m/z* 100-1700, spray voltage 4k V, drying gas flow rate 6 L/min, gas temperature 250°C, in full scan MS1 mode. The instrument was tuned and calibrated prior to each run. Agilent .d raw files were converted to mzML using ProteoWizard MSConvert software. Peaks were then aligned, grouped, and integrated using the xcms R package (BioConducter version 3.16). Isotope and adduct assignments were made using the R package CAMERA (BioConducter version 3.16). For lipidomics experiments in ACSL4 Control and KO cells, 3832 peaks were detected after xcms analysis. The AUC/mg protein levels of 1546 peaks were significantly changed ± palbociclib treatment in Control cells, as determined by two-tailed Student’s t-test (*P* < 0.05). 443 remained significant after Benjamini-Hochberg FDR testing with a strict cutoff of q < 0.01. Of those, 313 peaks were not altered by palbociclib treatment in ACSL4^KO1/2^ cell lines (two-tailed Student’s t-test, p < 0.05). Values are reported in the Table S2.

### Targeted MS/MS verification

Select lipids from the *MBOAT1* overexpression dataset and the ACSL4^KO^ dataset were validated by targeted MS/MS run on an Agilent 6545 qTOF. MS analysis was performed in positive mode and negative mode. Chromatography and instrument parameters were identical to those used for the untargeted lipidomics (see above) with the following additional settings for MS2. For all lipids, a collision energy of 20 eV was used and the isolation window was set to narrow (∼1.3 *m/z*). For identification of PE P-16:0/20:4 and PE P-16:0/22:6, 30,000 HT-1080^N^ cells were seeded in 6-well plates for DMSO treatment, and or 70,000 cells were seeded for palbociclib (2 μM) treatment. The next day, growth media were replaced with 2 mL of media containing either DMSO or palbociclib. 48 h later, lipids were extracted as described above for untargeted lipidomic analysis. 10 µL of lipid extract was run on the qTOF. In negative mode, analytes with *m/z* of 722.5127 and 746.5127 were fragmented. In positive mode, *m/z* 724.5273 and *m/z* 748.5273 were fragmented and analyzed. Only one species with *m/z* 722.5127 that was fragmented also displayed an increased abundance in HT-1080^N^ cells treated with palbociclib compared to those treated with DMSO. The same was true of *m/z* 746.5127. The MS/MS spectra of these two ions was analyzed in detail. In negative mode, the *sn*-2 chains of plasmalogen PE lipids can be identified as described.^59, 60^ Based on the product ions detected, as well as their relative abundances, these two species could be conclusively identified as PE P-16:0/20:4 and PE P-16:0/22:6, as previously described (https://www.biorxiv.org/content/10.1101/2021.01.20.427444v1.full.pdf).

For identification of *m/z* 700.5287, 30,000 HT-1080^N^ rtTA-MBOAT1^WT^ and HT-1080^N^ rtTA-MBOAT1^H381A^ mutant cells were seeded into one well of a 6-well plate. The next day, cells were treated with either Dox (0.1 μg/mL) or an equivalent volume of sterile H_2_O. 48 h later, lipids were extracted as described above. 10 µL of lipid solution was run on the qTOF as described above. In negative mode, species with *m/z* 700.5287 were fragmented and analyzed. In positive mode, species with *m/z* of 702.543 were fragmented and analyzed. Species were selected for further analysis if their abundance increased in HT-1080^N^ rtTA-MBOAT1^WT^ cells treated with Dox, but not in HT-1080^N^ rtTA-MBOAT1^H381A^ cells treated with Dox. In negative mode, one species met these criteria. In positive mode no interpretable spectra for species with *m/z* 702.543 could be obtained. Therefore, the identity of this species could not be exclusively assigned. However, the results from the negative mode fragmentation indicated that this species was a PE phospholipid with 1-*O*-alkyl 16:1 on its *sn*-1 chain and 18:1 on its *sn*-2 chain. It was therefore annotated PE *a-*16:1/18:1 where *a* denotes the alkyl bond. For all MS/MS experiments, Agilent .d files were analyzed using Agilent MassHunter Qualitative Analysis software.

### Untargeted lipidomics in EMP2 Control and KO cells

On day 0, 300,000 HT-1080^N^ EMP2^WT^ or HT-1080^N^ EMP2^KO^ cells were seeded in 10 cm^2^ plates for DMSO treatment. For palbociclib treatment, 600,000 cells were seeded. On day 1, 2 μM palbociclib (or an equal volume of DMSO) was added to the growth medium. 48 h later (day 4), cells were washed with 3 mL of 4°C 1X PBS and scraped into 0.5 mL of cold PBS. Cells were pelleted at 450 x *g* for 5 min, supernatant was aspirated, and pellets were stored at −80°C. Before extraction, cell pellets were thawed on ice for 30 min and resuspended in 50 μL PBS. 3 μL of SPLASH® LIPIDOMIX® internal standard (Cat# 330707-1EA, Avanti Polar Lipids) dissolved in methanol was added directly to each suspension. Lipids were extracted by adding 1.25 mL of tert-butyl methl ether (Cat# 34875-1L, Sigma Aldrich) and 0.375 mL of methanol (Cat# A452-4, Fisher Scientific). The mixture was incubated on an orbital mixer for 1 h (room temperature, 32 rpm). To induce phase separation, 0.315 mL of H_2_O was added, and the mixture was incubated on an orbital mixer for 10 min (room temperature, 32 rpm). Samples were centrifuged (room temperature, 2 min, 15,000 x *g*). Upper organic phase with collected and subsequently dried *in vacuo* (Eppendorf concentrator 5301, 1 ppm).

Dried lipid extracts were reconstituted in chloroform/methanol (150 μL, 2:1, v/v) and 20 μL of each extract was transferred to HPLC vials containing glass inserts. Quality control samples were generated by mixing equal volumes of each lipid extract followed by aliquotation in 20 μL aliquots. Aliquoted extracts were dried *in vacuo* (Eppendorf concentrator 5301, 1 ppm) and redissolved in 20 μL of 2-propanol (Cat# 1.02781.4000, Supelco, St. Louis, MO) for injection.

Lipids were separated by reversed phase liquid chromatography on a Vanquish Core (Thermo Fisher Scientific, Bremen, Germany) equipped with an Accucore C30 column (150 x 2.1 mm; 2.6 µm, 150 Å, Thermo Fisher Scientific, Bremen, Germany). Lipids were separated by gradient elution with solvent A (MeCN (Cat# A955-4, Fisher Chemical)/H_2_O (Cat# W6-4, Fisher Chemical), 1:1, v/v) and B (2-propanol/MeCN/H_2_O, 85:10:5, v/v) both containing 5 mM NH_4_HCO_2_ (Cat# 70221-25G-F, Sigma-Aldrich) and 0.1% (v/v) formic acid (Cat# A117-50, Fischer Chemical). Separation was performed at 50°C with a flow rate of 0.3 mL/min using the following gradient: 0-15 min – 25 to 86 % B (curve 5), 15-21 min – 86 to 100 % B (curve 5), 21-32 min – 100 % B isocratic, 32-32.1 min – 100 to 25 % B (curve 5), followed by 5.9 min re-equilibration at 25 % B.

Reversed phase liquid chromatography was coupled on-line to a Q Exactive Plus Hybrid Quadrupole Orbitrap mass spectrometer (Thermo Fisher Scientific, Bremen, Germany) equipped with a HESI probe. Mass spectra were acquired in positive and negative modes with the following ESI parameters: sheath gas – 40 L/min, auxiliary gas – 10 L/min, sweep gas – 1 L/min, spray voltage – 3.5 kV (positive ion mode); −2.5 kV (negative ion mode), capillary temperature – 250 °C, S-lens RF level – 35 and aux gas heater temperature – 370 °C.

Data acquisition for lipid identification was performed in quality control samples by acquiring data in data dependent acquisition mode (DDA). DDA parameters featured a survey scan resolution of 140,000 (at *m/z* 200), AGC target 1e6 Maximum injection time 100 ms in a scan range of *m/z* 240-1200. Data dependent MS/MS scans were acquired with a resolution of 17,500, AGC target 1e5, Maximum injection time 60 ms, loop count 15, isolation window 1.2 *m/z* and stepped normalized collision energies of 10, 20 and 30 %. A data dependent MS2 was triggered when an AGC target of 2e2 was reached followed by a Dynamic Exclusion for 10 s. All isotopes and charge states > 1 were excluded. All data was acquired in profile mode.

For deep lipidome profiling, iterative exclusion was performed using the IE omics R package.^61^ This package generates a list for already fragmented precursors from a prior DDA run that can be excluded from subsequent DDA runs ensuring a higher number of unique MS/MS spectra for deep lipidome profiling. After the initial DDA analysis of a quality control sample, another quality control sample was measured but excluding all previously fragmentated precursor ions. Parameters for generating exclusion lists from previous runs were – RT window = 0.3; noiseCount = 15; MZWindow = 0.02 and MaxRT = 36 min. This workflow was performed one time to achieve a total of two DDA analyses of a quality control sample in positive and two DDA analysis in negative ionization mode. Data for lipid quantification was acquired in *Full MS* mode with following parameters – scan resolution of 140,000 (at *m/z* 200), AGC target 1e6 Maximum injection time 100 ms in a scan range of *m/z* 240-1200.

Lipostar (version 1.0.6, Molecular Discovery, Hertfordshire, UK) equipped with in house generated structure database featuring fatty acids with no information on double bond regio-or stereoisomerism covering glycerolipid, glycerophospholipid, sphingolipid and sterol ester lipid classes. The raw files were imported directly with a Sample MS Signal Filter Signal Threshold = 1000 for MS and a Sample MS/MS Signal Filter Signal Threshold = 10. Automatic peak picking was performed with an m/z tolerance = 5 ppm, chromatography filtering threshold = 0.97, MS filtering threshold = 0.97, Signal filtering threshold = 0.

Peaks smoothing was performed using the Savitzky-Golay smoothing algorithm with a window size = 3, degree = 2 and multi-pass iterations = 3. Isotopes were clustered using a m/z tolerance = 5 ppm, RT tolerance = 0.25 min, abundance Dev = 40%, max charge = 1. Peak alignment between samples using an m/z tolerance = 5 ppm and an RT tolerance = 0.25 min. A gap filler with an RT tolerance = 0.05 min and a signal filtering threshold = 0 with an anti-Spike filter was applied.

For lipid identification, a “MS/MS only” filter was applied to keep only features with MS/MS spectra for identification. Triacylgylcerols, diacylglycerols and sterol esters were identified as [M+NH4]+ adducts. Lysophosphatidylcholines, lysophosphatidylethanolamines, Acyl-, ether-and vinyl ether-PE, ceramides and Sphingomyelins were analyzed as [M+H]+ adducts. Phosphatidylserines, and phosphatidylinositols were analyzed as [M-H]-adducts. Acyl-, ether- and vinyl ether-Phosphatidylcholines were identified as [M+HCOO]-adducts. The following parameters were used for lipid identification: 5 ppm precursor ion mass tolerance and 20 ppm product ion mass tolerance. Automatic approval was performed to keep structures with quality of 3-4 stars. Identifications were refined using manual curation and Kendrick mass defect analysis and lipids that were not following these retention time rules were excluded as false positives. Quantification was performed by peak integration of the extracted ion chromatograms of single lipid adducts of these high confidence lipids. Peak integration was manually curated and adjusted. Identified lipids were normalized to peak areas of added internal standards to decrease analytical variation and eventually normalized to protein concentrations of cell pellets after lipid extraction.

### Statistical analysis of EMP2^KO^ lipidomics

678 total lipids were detected. Two-tailed t-tests followed by Benjamini-Hochberg FDR testing was performed comparing HT-1080^N^ Control to EMP2^KO^ cells, with an FDR cutoff of q < 0.01. When comparing DMSO to palbociclib treatment in Control cells, 2-tailed t-tests followed by Benjamini-Hochberg FDR testing was performed. No lipids were significant with an FDR cutoff of q < 0.01 and so an FDR cutoff of q < 0.05 was used instead. Lipids were included if palbociclib treatment and EMP2^KO^ altered the lipid in the same direction (positive vs negative). Lipids were included in the final count if they were not significantly different in the DMSO and palbociclib conditions in EMP2^KO^ cells (2-tailed t-test, p > 0.05). Lipid pathway enrichment was assessed using the Bioinformatics Methodology For Pathway Analysis (BioPAN) webtool (https://www.lipidmaps.org/biopan/). Networks were reported that contained >2 nodes. Principal component analysis was performed using MetaboAnalyst 5.0.^62^ Data was uploaded in .csv format to the Statistical Analysis [one factor] tool. Data was transformed using the “Log transformation (base 10)” setting and then scaled using the “Auto Scaling (mean-centered and divided by the standard deviation of each variable)” setting. PCA scores and loadings data were exported in .csv format and graphed using GraphPad Prism.

### Analysis of CTRP dataset

Data from the Cancer Therapeutics Response Portal (CTRP) v2.1 dataset (https://ocg.cancer.gov/ctd2-data-project/broad-institute-screening-dependencies-cancer-cell-lines-using-small-molecules-0) was accessed on June 12^th^, 2023. The data for statistically significant correlations between basal gene expression and small-molecule sensitivity across all cancer cell lines were extracted from the v21.data.gex_global_analysis.txt table and plotted using GraphPad Prism.

### In vivo administration of palbociclib

NOD-*scid* IL2Rgamma^null^ (NSG) mice were obtained commercially (Jackson Laboratory, stock no. 005557, Bar Harbor, ME). HT-1080^N^ tumor cells were mixed with Matrigel (Cat# 356231, Corning) in a 1:1 ratio such that 2.5 x 10^6^ cells in 200 µL of media was mixed with 200 µL of Matrigel. Tumors were injected into both the left and right flanks of mice and allowed to incubate until reaching an average volume of 100 mm^3^ across all tumors. Mice were then divided into cohorts that received either vehicle control (methocellulose grade A4M 0.5% w/v in milliQ water), or palbociclib (Cat# P-7788, LC Laboratories, Woburn, MA) dissolved in vehicle at 25 mg/mL. In the first experiment, mice were dosed at 100 mg/kg daily for 5 d by oral gavage. Mice were sacrificed on day 6 according to IACUC regulations. Tumors were extracted and dissociated for analysis of mRNA and lipid expression, which were performed as above. We were unable to successfully extract assay-quality mRNA for two tumors, and so mRNA from nine tumors (vehicle) and seven tumors (palbociclib) were analyzed in downstream analyses.

### In vivo combination of palbociclib and compound 28

NOD-*scid* IL2Rgamma^null^ (NSG) mice were obtained from the Jackson Laboratory (Stock No. 005557, Bar Harbor, ME). HT-1080^N^ tumor cells were mixed with Matrigel (Cat# 356231, Corning) in a 1:1 ratio such that 2.5 x 10^6^ cells in 200 µL of media was mixed with 200 µL of Matrigel. Tumors were injected into both the left and right flanks of mice and allowed to incubate until reaching an average volume of 100 mm^3^ across all tumors. Mice were then divided into cohorts that received either vehicle control (methocellulose grade A4M 0.5% w/v in milliQ water), or palbociclib (Cat# P-7788, LC Laboratories, Woburn, MA) dissolved in vehicle at 25 mg/mL. In a first experiment with palbociclib or vehicle alone, mice were dosed at 100 mg/kg daily for 5 d by oral gavage. Mice were sacrificed on day 6 according to IACUC regulations. Tumors were extracted and dissociated for analysis of mRNA and lipid expression.

For experiments testing the palbociclib and compound 28 combination, the vehicle for palbociclib was methocellulose as described above. For compound 28, a solution of methocellulose (0.5% w/v, Cat# ME137-100GM, Spectrum Chemicals, West Compton, CA) with Tween80 (0.5% w/v, Cat# P1754-500ML, Millipore Sigma, Burlington, MA) dissolved in ddH_2_O was used as the vehicle. This solution was termed MCT. Polyethylene glycol 400 (PEG400, Cat# PO110-500MLGL, Spectrum Chemical) and ethanol (Cat. 493546-1L, Sigma Aldrich) were combined in a 1:3 ethanol:PEG400 ratio (ethanol/PEG400). The final vehicle for compound 28 contained 40:60 ethanol/PEG400:MCT, and compound 28 was dissolved at 15 mg/mL. Female 3-month-old NSG mice (Stock No. 005557, Jackson Laboratories) were injected with HT-1080 cells as above, and the experiment began when tumors reached an average volume of 100 mm^3^ (day 0). Mice were dosed with compound 28 (60 mg/kg) and/or palbociclib (100 mg/kg) on the following schedule. Palbociclib or vehicle control was dosed on days 2, 3, 9 and 10. Compound 28 or appropriate vehicle control was dosed on days 5, 6, 12, and 13. Animals began to reach humane endpoints and so mice were sacrificed and tissues harvested on day 17. Analysis of lipid levels was performed as described above.

### Immunohistochemistry for 4-HNE

Isolated tumor xenografts were fixed in 10% neutral buffered formalin. Cassettes were then submerged in 70% ethanol for 24 h. Subsequent sample preparation and staining was performed at the Stanford Animal Histology core facility. Paraffin embedding was performed by submerging samples in 70% ethanol (40 min, 40°C), 95% ethanol (2X 40 min, 40°C), 100% ethanol (2x 40 min, 40°C), xylene (3x 40 min, 40°C), and subsequently submerged in paraffin (2x 40 min, 60°C). Samples were cut to a thickness of approximately 5 microns and affixed onto glass slides (UNSPSC Code 41110000, Springside Scientific LLC, Durham, NC). Once dry, the tissue slices were deparaffinized and antigen retrieval was performed by boiling samples in citric acid (pH 6.0) for 20 min. Samples were washed in TBS for 1 min and blocked in avidin blocking solution (Cat# 927301, BioLegend, San Diego, CA) for 10 min. Slides were washed with TBS for 1 min and then blocked in biotin blocking solution (Cat# 927301, BioLegend) for 10 min. Finally, slides were washed for 1 min with TBS and then blocked in 3% goat serum (gift from Stanford Veterinary Service Center) in TBS for 10 min. Slides were then incubated overnight at 4°C with anti-4-hydroxynonenol (4-HNE) primary antibody (Cat# ab46545, AbCam) at a concentration of 1:300 in 3% goat serum in TBS with 0.5% Tween. The next day, slides were washed in TBS for one minute, blocked with 3% Peroxide block (Cat# H312-500, Thermo Fisher Scientific) diluted 1:10 in PBS for 10 min, and then washed three times with TBS for three min. Ready to use (no dilution) BT goat anti-rabbit secondary antibody (Cat# ab64256, AbCam) was added to the slides for 45 min at room temperature. 3x TBS washes (3 min each) were performed, and then slides were incubated with streptavidin-HRP tertiary antibody (Cat# N100, Thermo Fisher Scientific) at 1:1,000 dilution for 45 min at RT. Slides were washed three times with TBS for 3 min each. DAB chromogen (Cat# ab64238, AbCam) was added to the slides for 5 min, and slides were washed in deionized water for 1 min. Slides were imaged on a Lionheart FX microscope (Agilent BioTek). Ten images were captured per slide. Representative images are displayed.

### Quantification of 4-HNE stain

A python script was developed to quantify the intensity of brown (4-HNE positive) pixels in each image. Python version 3.10 was used and code was developed and run in PyCharm version 2021.1.1. Briefly, RGB images were converted to YCoCg color space, as the Co vector was found to be a good representation of the staining data. The Co pixel data (default 0 to 255) was normalized from 0 to 1 (subsequently termed “intensity value”) by dividing all values by 255. Pixels were called as 4-HNE positive (4HNE+) if they met a minimum intensity threshold determined empirically. Flood fill,^63^ a background subtraction strategy, was used to remove background pixels from the analysis. The intensity values of 4HNE+ pixels were averaged to create a single intensity score per image. The intensity scores across all 380 images were normalized and plotted. Statistical testing was performed comparing the groups using two-tailed Student’s t-tests.

### Quantification, statistical analysis, and software

Lethal fraction calculations and statistical analyses were performed using Microsoft Excel 14.6.0 (Microsoft Corporation, Redmond, WA). Flow cytometry data were analyzed using FlowJo 10.6.1 (FlowJo LLC, Ashland, OR). C11 BODIPY 581/591 images acquired using the Lionheart imager were analyzed with Gen5 v3.10 software (BioTek). Graphing was performed using Prism 9 (GraphPad Software, La Jolla, CA). Figures were assembled using Adobe Illustrator (Adobe Systems, San Jose, CA). Details of experiment design and statistical testing (where appropriate) used can be found in the main text, figure legends, and Methods.

## Data availability

Underlying deviation scores for the nutlin-3 modulatory screen is available in Table S1. Processed data for lipidomics experiments are available in Table S2 and Table S5. Raw data for RNA-sequencing is available online at GEO (GSE180265), and processed data is available in Table S3. Uncropped immunoblots and new code will be made available online.

---

**Figure S1.**
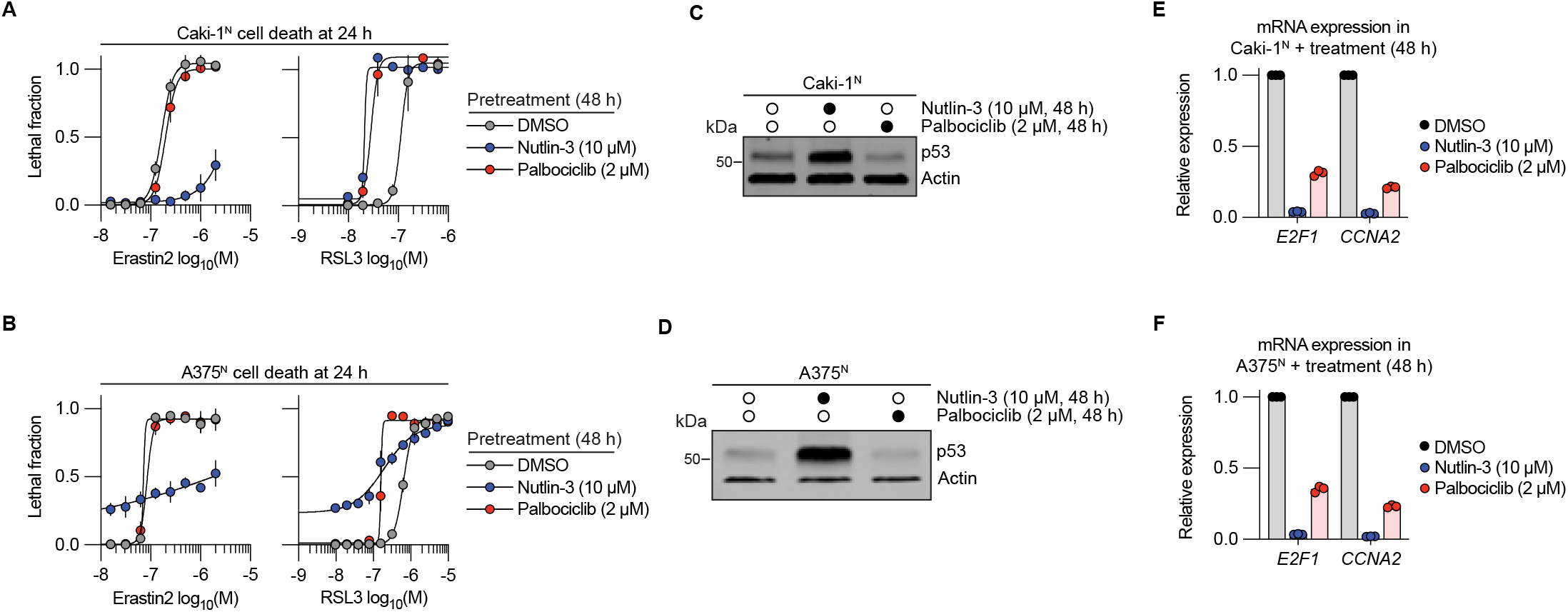
CDK inhibition sensitizes to ferroptosis. (A,B) Cell death measured by imaging following pretreatment. Note that for A375^N^ cells in (B) that nutlin-3 treatment alone causes a degree of baseline cell death. Results in represent mean ± SD from three independent experiments. (C,D) Protein abundance determined by immunoblotting. Blots are representative of three experiments. (E,F) Relative mRNA expression determined by reverse transcription and quantitive polymerase chain reaction (RT-qPCR) analysis. Individual datapoints from independent experiments are shown.

**Figure S2.**
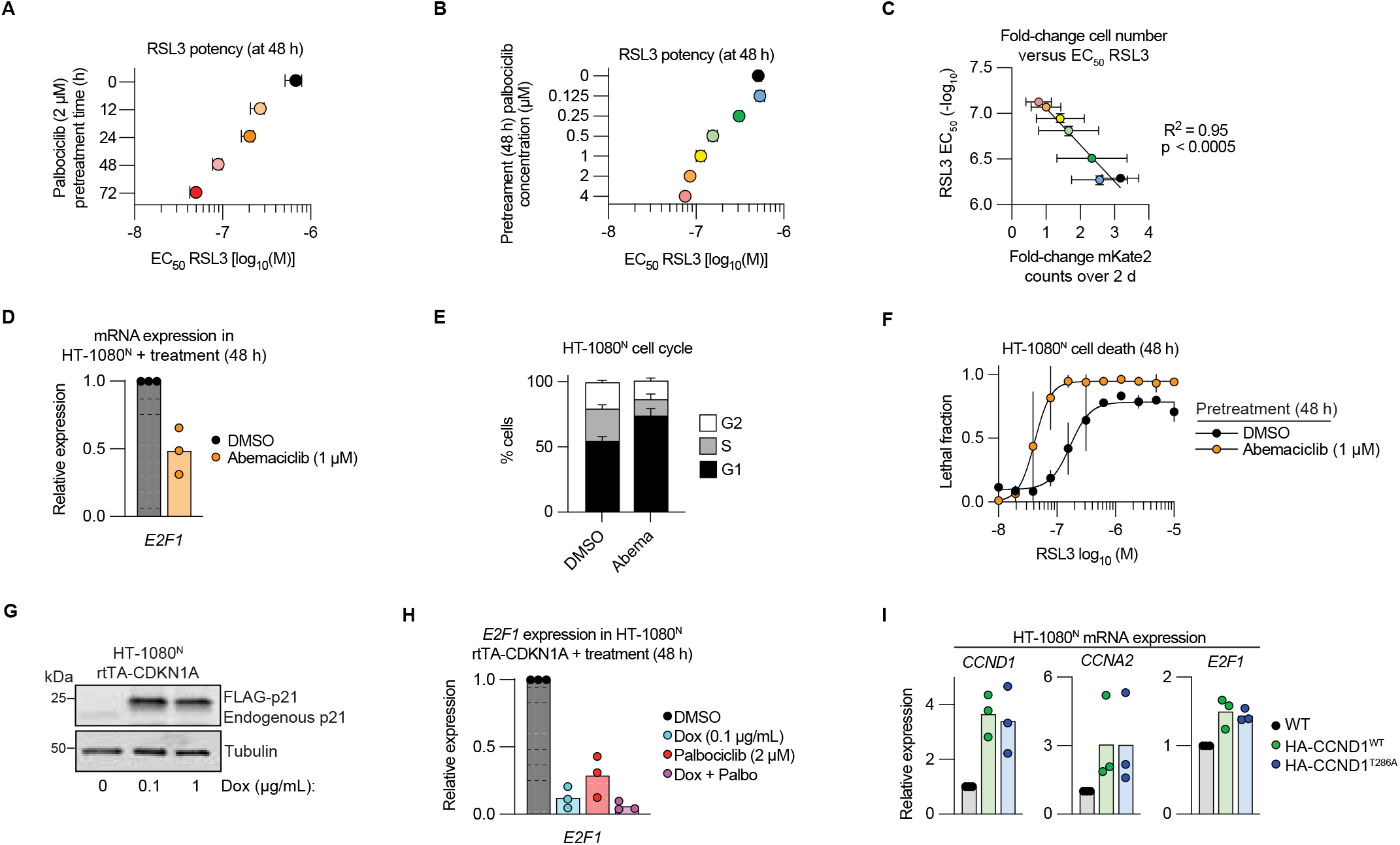
CDK inhibition sensitizes to ferroptosis. (A,B) RSL3 EC_50_ in HT-1080^N^ cells determined by imaging following various lengths of palbociclib pretreatment (A) or pretreatment with different concentrations of palbociclib for 48 h. Data repre-sent represent mean ± 95% confidence interval (CI). (C) HT-1080^N^ cell proliferation in response to treatment with palbociclib versus RSL3 EC_50_ determined in (B) following 48 h of palbociclib pretreatment. The color codes for individual datapoints correspond to those used in (B) and refer to the concentration of palbociclib used in the pretreatment. Pearson correlation values are reported. The data for proliferation is mean ± SD and the error for the RSL3 EC_50_ values are the 95% CI values from (B). (D) Relative gene expression determined by reverse transcription and quantitive polymerase chain reaction (RT-qPCR) analysis. (E) Cell cycle phase determined by propidium iodide (PI) staining and flow cytometry. Abema, abemaciclib (1 µM, 48 h). (F) Cell death measured by imaging following pretreatment. (G) Protein abundance determined by immunoblotting following doxycycline (Dox, 48 h) treatment. Blot is representative of three independent experiments. (H, I) Relative mRNA expression determined by RT-qPCR. Data in (E) and (F) are mean ± SD from three independent experiments. Individual data points from independent experiments are shown in (D), (H) and (I).

**Figure S3.**
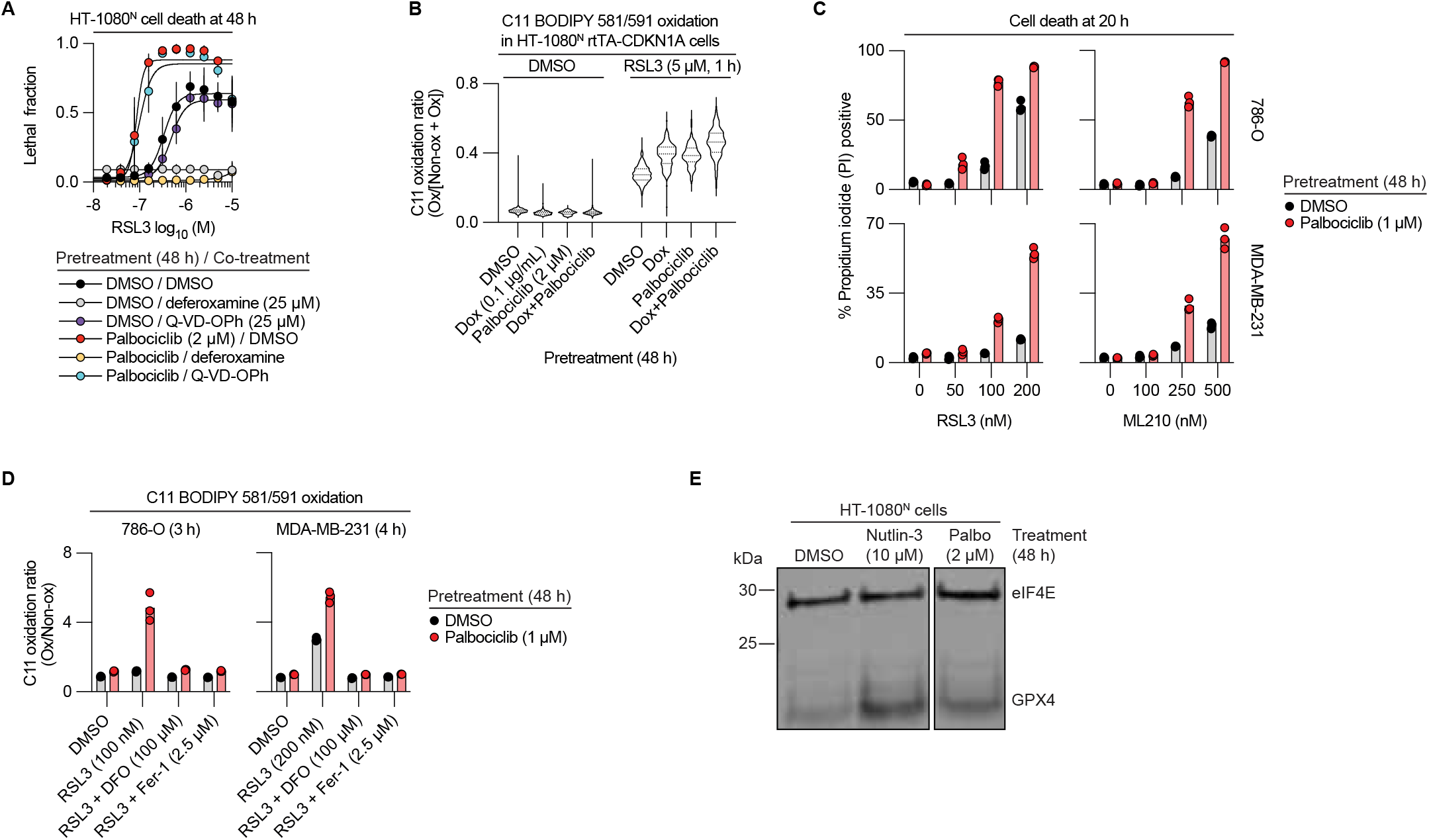
CDK inhibition enhances ferroptosis sensitivity. (A) Cell death measured by imaging following pretreatment. Data are mean ± SD from three independent experiments. (B) Lipid peroxidation detected using C11 BODIPY 581/591 (C11) and epifluorescent imaging in cells treated as indicated. Dox, doxycycline. Ox, oxidized. Non-ox, non-oxidized. Results are from 129-489 individual cells/condition. (C) Cell death measured by flow cytometry following pretreatment. (D) Lipid peroxidation detected using C11 and flow cytometry in cells treated as indicated. (E) Protein abundance determined by immunoblotting. Palbo, palbociclib. The blot is representative of three independent experiments.

**Figure S4.**
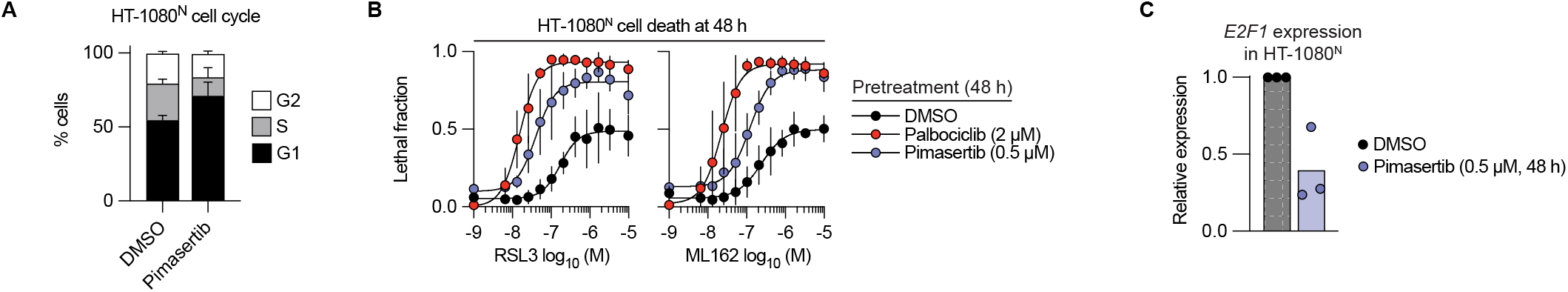
MEK1/2 inhibition enhances ferroptosis sensitivity. (*A*) Cell cycle phase determined by propidium iodide (PI) staining and flow cytometry. Pimasertib treatment was 0.5 µM, 48 h. (*B*) Cell death measured by imaging following pretreatment. (*C*) Relative mRNA expression determined by reverse transcription and quantitative polymerase chain reaction (RT-qPCR) analysis. Individual datapoints from independent experiments are shown. Results in (*A*) and (*B*) represent mean ± SD from three independent experiments.

**Figure S5.**
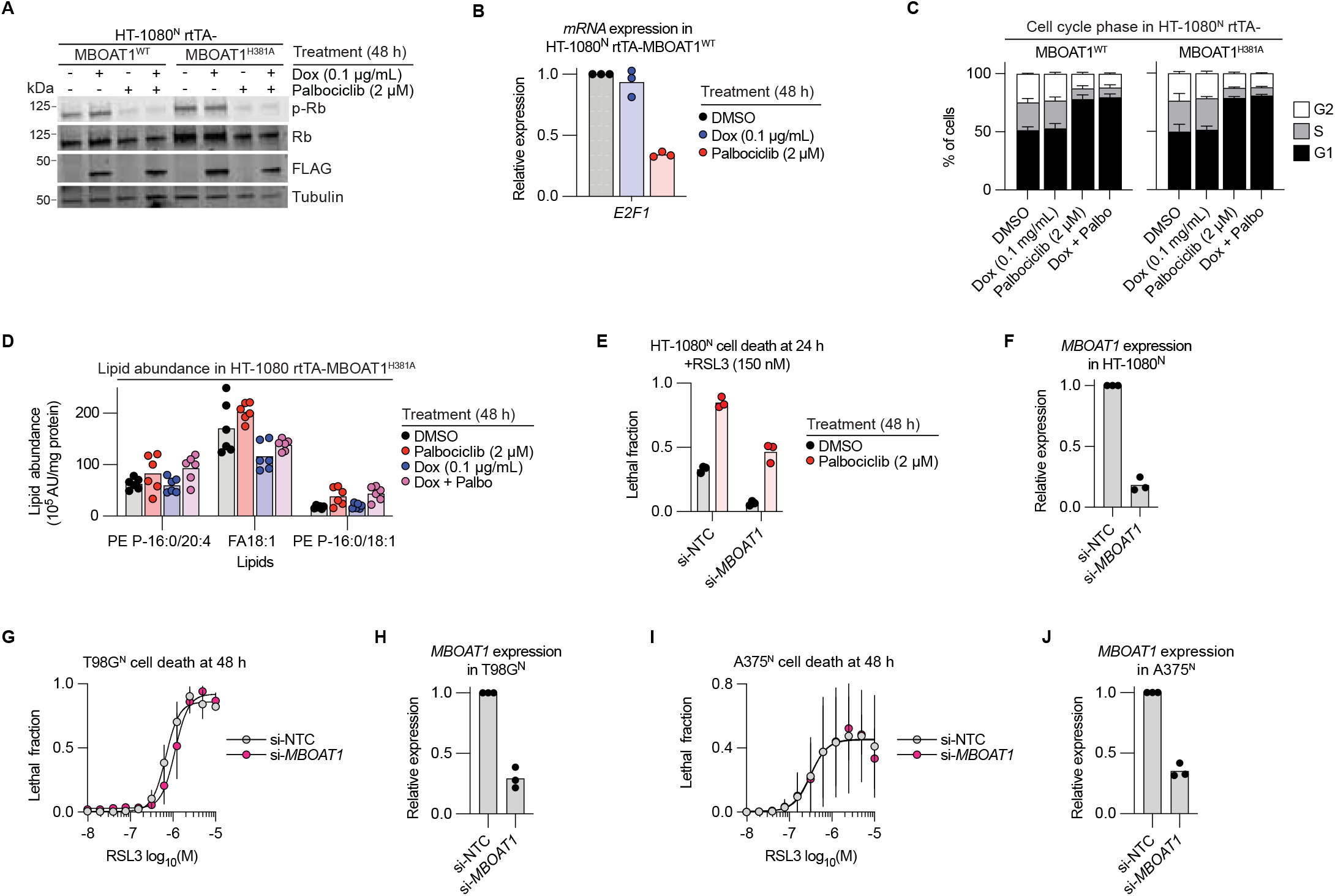
MBOAT1 and ferroptosis sensitivity. (A) Protein abundance determined by immunoblotting. Representative of three experiments. (B) Relative mRNA expression determined by RT-qPCR analysis. (C) Cell cycle phase determined by propidium iodide (PI) staining and flow cytometry. (D) Lipid abundance determined using liquid chromatography coupled to mass spectrometry. AU, arbitrary units. (E) Cell death determined by imaging. si, short interfering RNA. NTC, non-targeting control. Genetic silencing and DMSO or palbociclib treatments were performed at the same time for 48 h, prior to exposure to RSL3. (F) Relative mRNA expression determined by RT-qPCR analysis. (G) Cell death determined by imaging. (H) Relative mRNA expression determined by RT-qPCR analysis. (I) Cell death determined by imaging. (J) Relative mRNA expression determined by RT-qPCR analysis. Individu-al datapoints from independent experiments are shown in (B), (D-F), (H) and (J). Results in (C), (G) and (I) are mean ± SD from three independent experiments.

**Figure S6.**
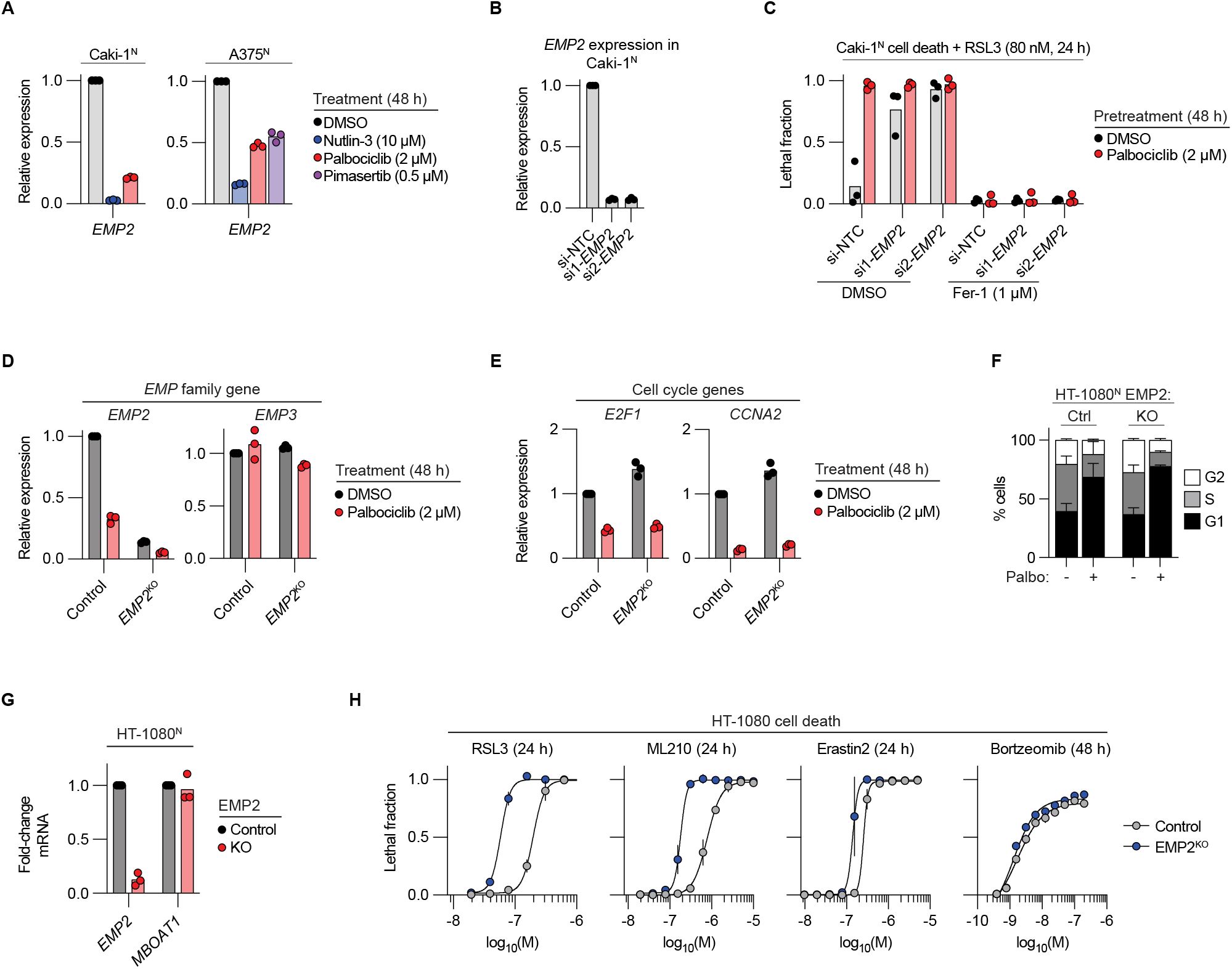
EMP2 and ferroptosis sensitivity. (A,B) Relative mRNA expression determined by reverse transcription and quantitative PCR (RT-qPCR) analysis. si, short interfering RNA. NTC, non-targeting control. (C) Cell death determined by imaging. (D,E) Relative mRNA expression determined by reverse transcription and quantitative PCR (RT-qPCR) analysis. (F) Cell cycle phase determined by propidium iodide (PI) staining and flow cytometry. Palbo, palbociclib (2 µM, 48 h). Ctrl, Control. (G) Relative mRNA expression determined by reverse transcription and quantitative PCR (RT-qPCR) analysis. (H) Cell death determined by imaging. Individual datapoints from independent experiments are shown in (A-E), (G). Data in (F), (H) represent mean ± SD from three independent experiments.

**Figure S7.**
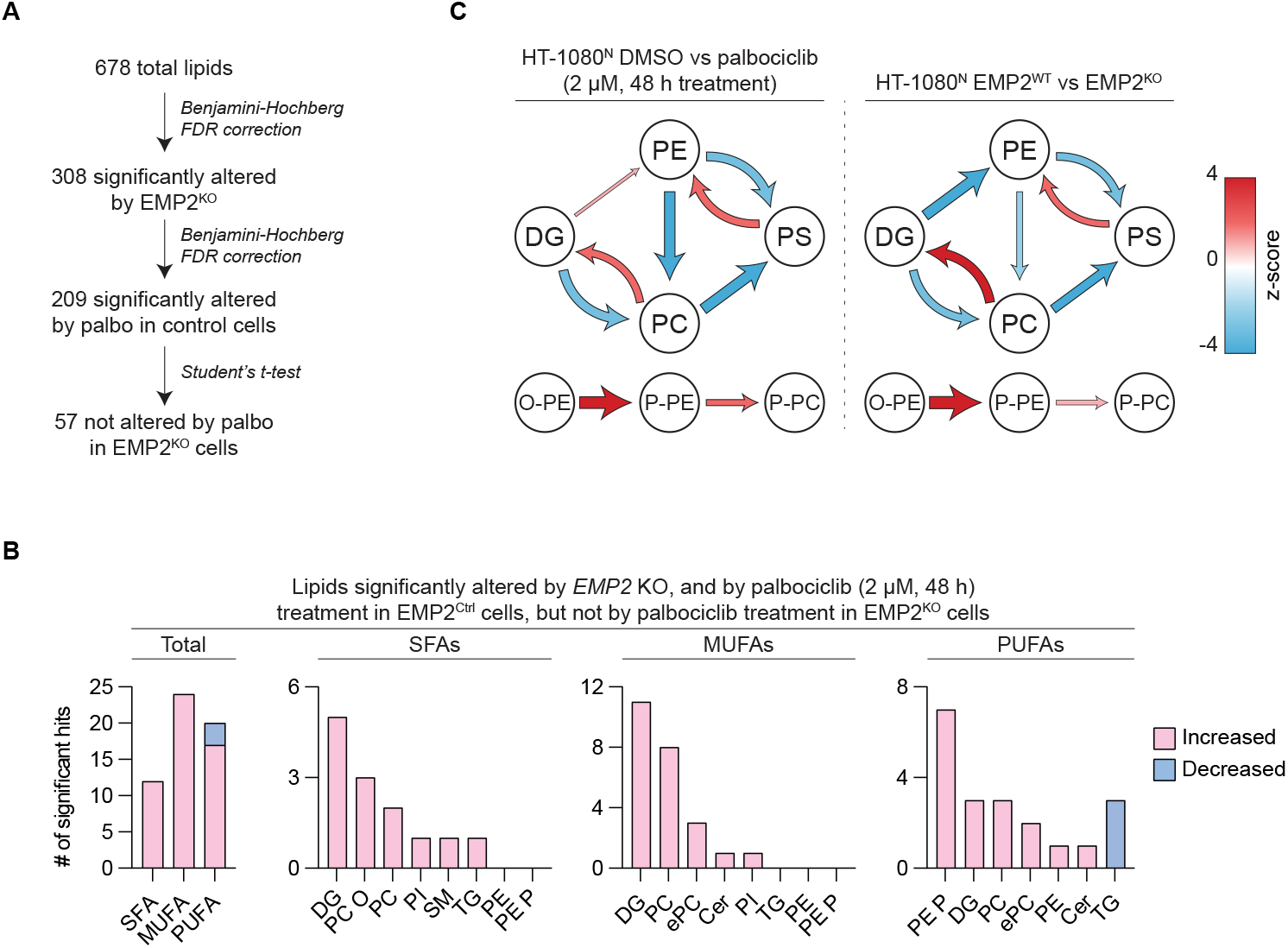
Lipidomic analysis of EMP2 gene disruption. (A) Strategy for the lipidomic analysis. FDR q < 0.01 for Control vs EMP2^KO^ cells, FDR q < 0.05 for Control DMSO vs palbociclib (2 µM, 48 h) conditions, and p < 0.05 (2-tailed Student’s t-test) for EMP2^KO^ DMSO vs palbociclib (2 µM, 48 h) conditions. (B) Analysis of 57 significantly altered lipids following statistical filtering. SFA, lipids that contained only saturated fatty acids. MUFAs, lipids that contained at least one monounsaturated fatty acid and no polyunsaturated fatty acids. PUFAs, lipids that contained at least one polyunsaturated fatty acid. DG, diglyceride. Cer, ceramides. PC, phosphatidylcholine, PC O, phosphatidylcholine ether lipids. ePC, phosphatidylcholine ether and plasmalogen lipids. PE, phosphatidylethanolamine. PE P, phosphatidylethanolamine plasmalogen. PI, phosphatidylinositol. SM, sphingomyelin. TG, triglyceride. (C) A summarized network of lipidomic changes derived from Bioinformatics Methodology for Pathway Analysis (BioPAN). Arrows are sized to exact z-scores.

